# Probing the zooarchaeological record across time and space for ancient pathogens

**DOI:** 10.1101/2025.02.12.637901

**Authors:** Anne Kathrine W. Runge, Ian Light-Maka, Ken Massy, Marcel Keller, Simon Trixl, Helja Kabral, Casey L. Kirkpatrick, Kirsten Bos, Jana Eger, Michal Ernée, René Kyselý, Michael Hochmuth, Dominik Poradowski, Aleksander Chrószcz, Norbert Benecke, David Daněček, Jana Klementová, Anatoli Nagler, Alexey A. Kalmykov, Anatoly R. Kantorovich, Vladimir E. Maslov, Andrey B. Belinskiy, Christiana L. Scheib, Meda Toderaş, Svend Hansen, Philipp W. Stockhammer, Kai Kaniuth, Regina Uhl, Sabine Reinhold, Rosalind E. Gillis, Elizabeth A. Nelson, Kamilla Pawlowska, Felix M. Key

## Abstract

Zoonoses are among the greatest threats to human health, with many zoonotic pathogens believed to have emerged during prehistory. Palaeomicrobiological investigations of the zooarchaeological record hold potential to uncover the reservoirs, host ranges, and host adaptations of zoonotic pathogens but face challenges in identifying promising specimens and pathogen DNA preservation. We performed palaeopathological and genetic examinations of 346 skeletal elements from domesticated and wild animals collected from 34 Eurasian sites dating across the last six millennia. We identified 68 signatures of ancient (opportunistic) pathogens, including the important zoonotic pathogen *Salmonella enterica*, and found support that palaeopathological lesions provide guidance for specimen selection. For two pathogen species, *Erysipelothrix rhusiopathiae* and *Streptococcus lutetiensis*, we confirmed their ancient authenticity using phylogenetics, showcasing an approach to explore the relationship between ancient low-coverage genomes and their modern-day relatives. Our work presents a pathway to understanding prehistoric zoonotic diseases by integrating zooarchaeological, palaeopathological, and genetic data.

## Introduction

Infectious diseases are a major health concern causing an estimated 13.7 million deaths worldwide in 2019 ^1^. Approximately 60% of human pathogens and 75% of emerging diseases are believed to have originated from animals ^2^. Such zoonotic spillover events can have devastating consequences when novel pathogens enter naive populations as evidenced by the recent Covid-19 pandemic ^3^, but zoonoses are not a new phenomenon. For instance, the 1918 flu pandemic likely had an avian origin 4 5, while outbreaks of plague, which are transmitted from rodent hosts via flea vectors, led to the Justinianic plague in 541–543 CE ^6^ and the Black Death in 1346–1353 CE ^7^. Although Palaeolithic hunter-gatherers came into contact with diseased animals, many zoonotic diseases were likely introduced into human populations following the Neolithic Transition starting around 10,000 years ago ^8,9^. In particular, the intensification of inter-species interactions between humans and their domesticated animals during livestock management, butchering or consumption of animal-derived products increased the likelihood of zoonotic transmission events ^8^. Furthermore, the Neolithic period introduced complex changes in the spatial organisation of settlements, demographic expansion, and subsistence patterns, which favoured the introduction and persistence of crowd diseases ^10,11^. Today, this is recognised in the One Health concept arguing that human health is interconnected with animal and environmental health with consequences ranging from zoonoses, antimicrobial resistance to food safety ^12^. Ancient DNA holds promise to be a powerful tool for exploring the One Health concept in deep time - recently also coined One Palaeopathology ^13^ - by reconstructing pathogen genomes from the zooarcheological record, revealing prehistoric disease reservoirs, retracing pathogen distribution, and uncovering adaptations to the human host.

While screening for microbes in large genomic datasets generated for ancient human population studies has become reasonably routine, the same cannot be said for zooarchaeological datasets. As a result, limited information is available concerning disease reservoirs in ancient and historic animal populations. Nevertheless, recent advances provided some notable exceptions highlighting the power of pushing palaeomicrobiology into the zooarcheological record. A *Yersinia pestis* genome from Bronze Age domesticated sheep provided insights into the host range and evolution of a pathogen lineage thus far exclusively identified in human archeological remains ^14^. Furthermore, an 8,000 year old *Brucella melitensis* genome from sheep confirms the emergence of the zoonosis Brucellosis during the Neolithic Transition ^15^, a Medieval strain of *Mycobacterium leprae* confirmed squirrels as a long-term animal host of the pathogen ^16^, and reconstructed Marek disease virus genomes informed upon the origin and virulence of a contemporary livestock infection ^17^. Despite these promising initial results, no systematic investigation has been conducted to test the preservation of microbial pathogen genomes in zooarchaeological remains.

The detection of microbial pathogen DNA from animal remains excavated at human settlement sites is particularly challenging because they typically enter the archaeological record following human processing. This has multiple implications for pathogen DNA preservation. First, the majority of animals under human management died from slaughter rather than natural causes, including infectious diseases. In fact, an additional confounding factor would be the practice of culling sick animals, further reducing the likelihood of diseased specimens being preserved. Second, unlike human remains, which were more frequently intentionally buried, the majority of animal remains were discarded as household waste, resulting in prolonged exposure to the environment and thus degrading target DNA^18^. Finally, animal remains may have experienced amplified DNA degradation due to heating, boiling, or roasting during preparation for consumption ^18^. Altogether these factors likely decrease the chance of recovering pathogen DNA from animals, explaining why only a few studies have described pathogen genomes reconstructed from ancient DNA from animals. Nevertheless, in some cases, signs of disease are preserved within the archaeological record in the form of palaeopathological lesions ^19,20^. Such lesions typically occur as a result of prolonged disease, but while they are frequently recorded and studied in human remains, the same is rarely the case for animals ^19^. Once identified, however, they may allow researchers to overcome at least in part the challenges of palaeomicrobiology in the zooarcheological record.

In this study we investigate hundreds of zooarchaeological specimens from across Eurasia for pathogen DNA, with a particular focus on the Bronze Age. We use palaeopathology to identify lesions that may have resulted from infectious diseases. We use these lesions as targets for ancient DNA sampling to increase the probability of recovering ancient pathogen DNA and identify known pathogens that were present in prehistoric animal populations. Our results show ancient DNA signatures of ancient (opportunistic) pathogens within the zooarchaeological record and highlight the power of palaeopathology to prioritize specimens for pathogen recovery, opening a new direction in palaeomicrobiology.

## Results

### Zooarchaeological collection of 346 skeletal elements from across Eurasia

To test the preservation of microbial pathogen DNA within the zooarchaeological record we collected a total of 346 skeletal elements from 329 individual animals discovered at 34 archaeological sites across Eurasia (**Figure 1a**, **Table S1**). The majority of these sites are located in Europe, but two sites, Monjukli Depe and Tilla Bulak, are located in Central Asia. The sites span approximately 5,800 years of human history (**Figure 1b**), with the oldest site, Monjukli Depe, dated to 4650-4350 BCE and the youngest site, Giecz 10, dated to 900-1200 CE. The sites cover periods dating to the Neolithic to the Medieval period, with the majority of sites belonging to the Bronze Age. We focused on investigating Bronze Age zooarchaeological specimens because human-derived ancient zoonotic-pathogen genomes were repeatedly identified in specimens from the Bronze Age, a period of major human migratory events and the Neolithization in Eurasia ^21–26^.

**Figure 1.**
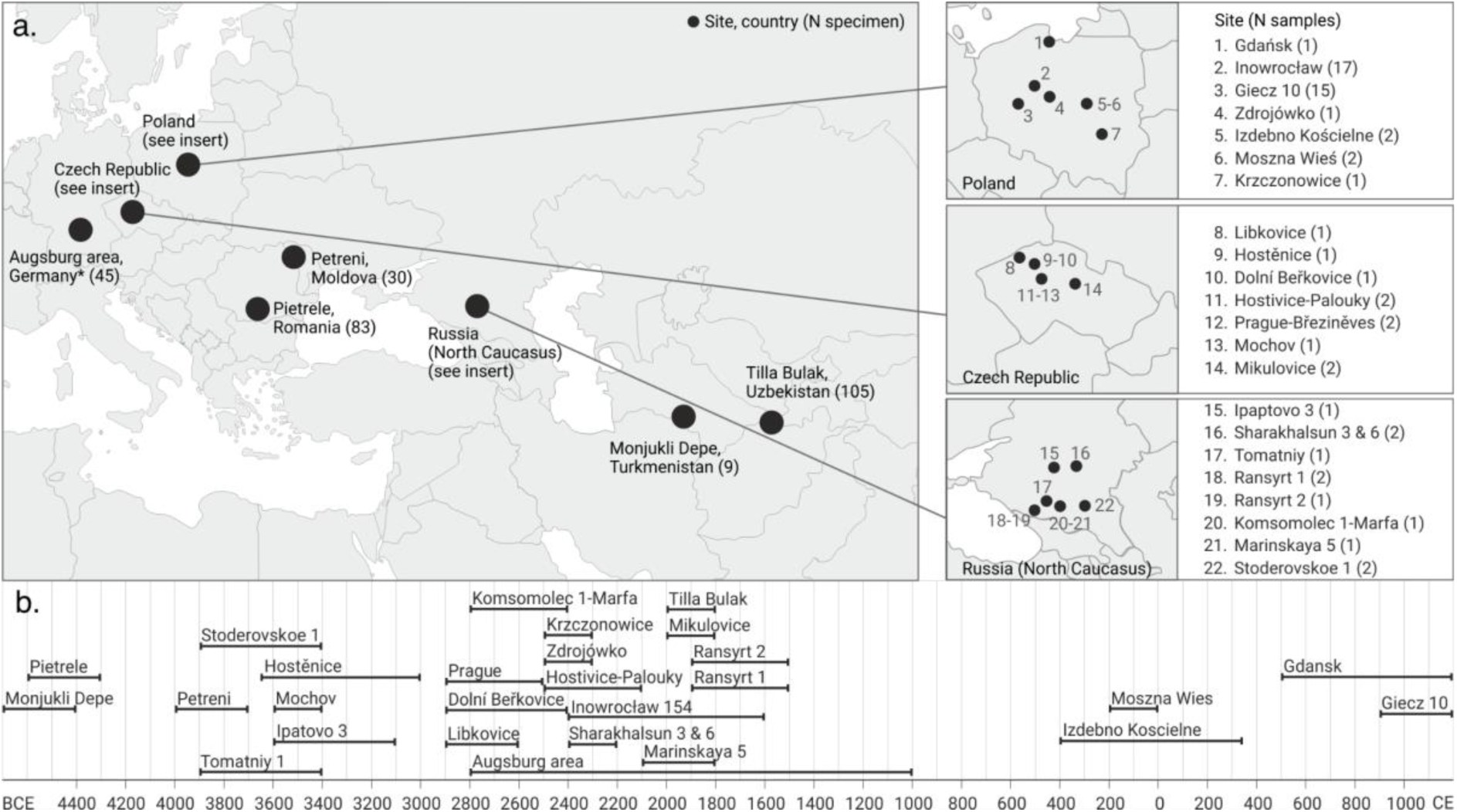
Geographical and temporal distribution of 346 zooarchaeological specimens. **a.** The 34 archaeological sites included in this study are distributed across Europe and Central Asia. Cut-out maps are included for countries with more than one site. The Augsburg area (Germany) consists of seven sites in and around Augsburg, which are not shown individually on the map. Coordinates are available in **Table S1**. **b.** The timeline shows the age ranges of each site included in this study, which are estimates based on C14 dates of associated specimen or archaeological dating based on features (**Table S1** and **Supplemental Online Material**). Note that the Bronze Age covers different absolute chronological horizons within different geographical locations.

The specimen selection was aimed primarily at domesticated species defined by morphology, but wild animals were also included, especially if their abundance indicated that they contributed significantly to the local economy. All specimens were taxonomically assigned based on morphology and later also classified using the ancient host DNA content of the generated metagenomic data (see below for data generation). In total, we collected 107 cattle, 99 sheep, 45 pigs/wild pigs, 18 dogs, and 12 goats along with 51 remains from species that were less abundant than those previously mentioned or could not be classified to a lower taxonomic unit (**Figure 2, Table S2**).

**Figure 2.**
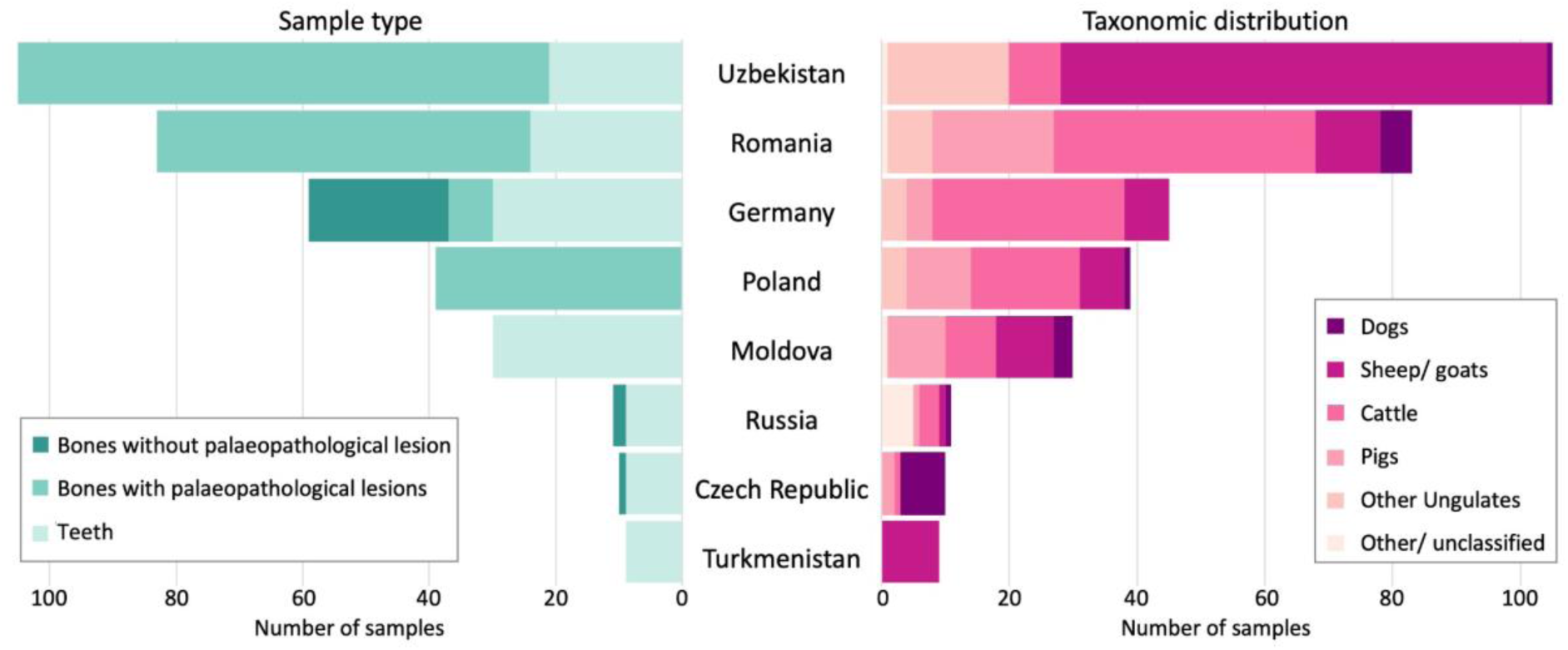
Sample type and taxonomic distribution across countries. Left: Showing the distribution of the 346 skeletal elements (bones with palaeopathological lesions, bones with no lesions, and teeth) that were sampled at each country. Right: Taxonomic distribution of the 329 sampled individual animals across countries illustrates that the majority of samples originated from domesticated species. The ‘other ungulates’ category includes ungulates other than sheep, goat, and cattle as well as ungulates that could not be classified at the species level. The ‘other/ unclassified’ category includes all other taxonomic groupings than those already mentioned. Note that there are fewer individuals than samples from Germany, because both teeth and bone were sampled from some individuals. See **Table S2** for detailed taxonomic classification of all samples.

### Palaeopathological and taphonomical evidence in the zooarchaeological record

In total, we collected 189 bones with palaeopathologies (54.6%), 25 bones with no identified palaeopathological changes (7.2%), and 132 teeth (38.2%) from the 34 sites included in this study. From one individual from Pietrele and seven of the German individuals, we collected both bones and one or more teeth. Bones from the sites in Germany, Poland, Pietrele (Romania), and Tilla Bulak (Uzbekistan) were visually inspected for palaeopathological lesions (**Figure 3, Table S2**). Although teeth can have lesions such as caries, none were observed in the studied material, and none of the teeth included in the study displayed pathological conditions. Identification of skeletal signs of infectious disease among the zooarchaeological specimens can be difficult to discern from unrelated environmental and taphonomic processes ^27^. The investigated specimens were deposited in sand, silt, and gypsum, and sufficiently well-preserved for palaeopathological analysis. We macroscopically screened the remains for various conditions that can be associated with, though not necessarily unique to, infection. This included: periostitis (inflammation of the periosteum, a connective tissue layer on the surface of bones, that is sometimes caused by infection), skeletal trauma (which may create entry sites for pathogens), lytic lesions and/or pathological new bone development (sometimes associated with infectious disease), alveolar recession (indicative of periodontitis, a severe gum infection), ante-mortem tooth loss (also sometimes associated with oral infections, such as periodontitis, caries and/or abscess), and arthropathy (sometimes related to infectious disease) ^19^.

**Figure 3:**
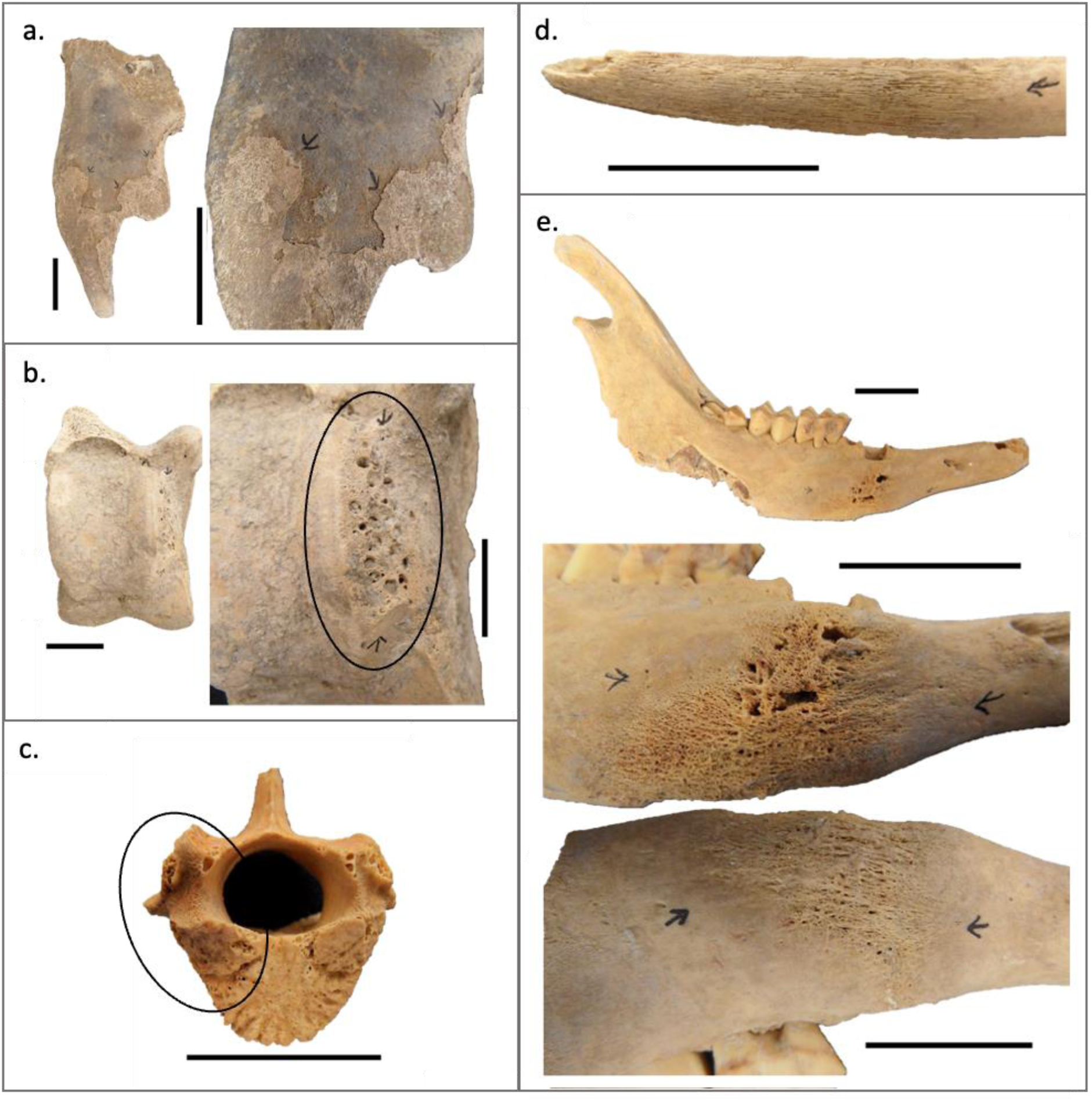
Examples of palaeopathologies that were sampled for this study and produced pathogen signatures in the screening. Arrows or circles indicate location of lesions. **a.** AZP-115, a piece of pig skull with woven periostitis, possibly associated with inflammatory diseases, **b.** AZP-183, a cattle astragalus with macroporosity suggestive of inflammation and degenerative joint disease possibly associated with infectious disease, **c.** AZP-268, a thoracic vertebra from a sheep with macroporosity on the right articular surface of the vertebral body and part of the neural arch (circled), possibly linked to inflammation or infection, **d.** AZP-195, a small ruminant rib with periostitis suggestive of infectious disease, **e.** AZP-222, a sheep mandible with marked active infection evidenced by reactive bone with abscess development. See **Table S2** for additional information. Scale bars indicate 2 cm.

Across all initially investigated sites only a minority of the zooarchaeological deposits showed any palaeopathological evidence but those included a variety of lesion types affecting a wide range of skeletal elements in many species (**Figure 3**, **Table S2**). More specifically, the bone assemblages from Poland, Romania (Pietrele), and Uzbekistan (Tilla Bulak) showed pathological lesions including periostitis, osteolytic lesions, oral pathologies, and arthropathy. Only some of the specimens from Germany (Augsburg area) displayed palaeopathological changes, including periostitis, osteomyelitis, and periodontitis, while the majority of collected specimens had no visible lesions. Across the investigated sites, a wide range of primarily domestic but also wild species is represented among the specimens with palaeopathological lesions including: horses, cattle, pigs, sheep, goats, dogs, red deer, aurochs, gazelles, and a fox and a beaver. Overall, sheep were dominant among the species showing palaeopathological lesions in our dataset (80 lesional specimens out of 192), but not significantly overrepresented after multiple testing corrections (Methods). Similarly, lesions were identified on a wide range of skeletal elements with mandibles (n=27), vertebrae (n=25), and ribs (n=24) being the majority. Collectively, the zooarchaeological specimen collection included a vast range of species and bone types covering different palaeopathological lesions suggestive of infection and thereby pathogens whose DNA fingerprint we can test.

### Identification and authentication of ancient pathogen signatures

From the 346 skeletal elements, we produced a total of 357 DNA extracts. These included 20 subsamples collected from 11 bones as well as nine teeth collected from four individuals (see **Table S2** for further details). Samples were sequenced on the Illumina platform, which, apart from five failed samples, generated between 2,324,302 and 90,823,912 sequencing reads per sample (**Table S2**). All sequencing reads which passed quality control thresholds and did not map to the human reference genome (see Methods) were taxonomically classified. The presence of DNA molecules from a range of bacterial, viral, and parasitic species was investigated (**Table S3**) using the HOPS pipeline ^28^ as implemented in nf-core/eager ^29^. We focus our analysis on species that are pathogenic to humans and animals, including species that are opportunistic pathogens able to colonise without causing disease in the host. The HOPS pipeline enables the identification of target DNA in a metagenomic dataset and at the same time interrogates signatures for its ancient authenticity, for example mismatches due to deamination at terminal bases of DNA fragments (ancient DNA damage) 30,31 or distribution of aligned reads along the genome. We applied stringent authentication criteria to ensure that any pathogens identified in the screening dataset were robust. These included a minimum of 50 reads aligning to the target taxonomic node, terminal bases showing a minimum of 10% DNA damage (for single-stranded libraries, 5’ or 3’ terminal C to T; for double-stranded libraries C to T at the 5’ terminal base or G to A at the 3’ terminal base), a declining edit distance, and little to no read stacking except for samples with exceptionally high number of reads (Methods).

From our screening dataset, 55 libraries (15%) produced high confidence bacterial hits that met our identification and authentication criteria (**Figure 4**). 31 samples produced hits to only one bacterial species, while 24 samples produced hits to more than one species. No hits to eukaryotic parasites or viruses were identified in our dataset. The authenticated bacterial species differed in their described pathogenic potential with some being identified as primarily pathogenic while others are opportunistic pathogens that can also asymptomatically colonise the host. The mostly pathogenic bacteria included the species *Salmonella enterica, Bordetella petrii*, *Coxiella burnetii*, and *Erysipelothrix rhusiopathiae* (**Figure 3**). *S. enterica* is a major zoonotic pathogen consisting of hundreds of serovars, with different host specificities of which some are known to infect humans causing gastroenteritis or systemic disease ^32,33^. Previously presented ancient human-derived *S. enterica* genomes from up to 6,000 years ago suggested host adaptation happened during the Neolithization period^22^; our results suggest that indeed animal-derived *S. enterica* genome signatures can be recovered to estimate its prehistoric host range, including sheep, goats, and dogs. *Bordetella* species are frequent pathogens of humans and animals, but the clinical relevance of *B. petrii* is unclear ^34^. *C. burnetii* is an obligate intracellular pathogen, which causes the disease coxiellosis in animals although infected individuals are often asymptomatic 35. *E. rhusiopathiae* infections have been characterised in a wide range of vertebrate and invertebrate species ^36^, and was recently identified in human remains from medieval Southwest Europe ^37^. It is the etiologic agent of swine erysipelas ^38^, but we identify it in cattle where it is known as an opportunistic pathogen and has been associated with septicemia, abscesses in the liver and lungs, and arthritis ^39^. The other identified bacterial species belong to the oral and gastrointestinal microbiome. These include *Corynebacterium stationis*, *Corynebacterium xerosis, Escherichia coli*, *Porphyromonas gingivalis*, *Veillonella parvula*, *Yersinia intermedia*, as well as several species of *Enterococcus*, *Staphylococcus* and *Streptococcus*. While the majority of these species are members of the stable microbiome of animals or, when information on animals was limited, of humans, most are known opportunistic pathogens that rarely cause infection in healthy individuals, but can cause disease in immunocompromised individuals. Among the identified oral microbiota we detect bacteria, incl. *P. gingivalis*, that are part of the so-called ‘red complex’ which are associated with severe forms of periodontal disease ^40^ and point to a reservoir of bacteria causing gum infections among humans and livestock.

**Figure 4.**
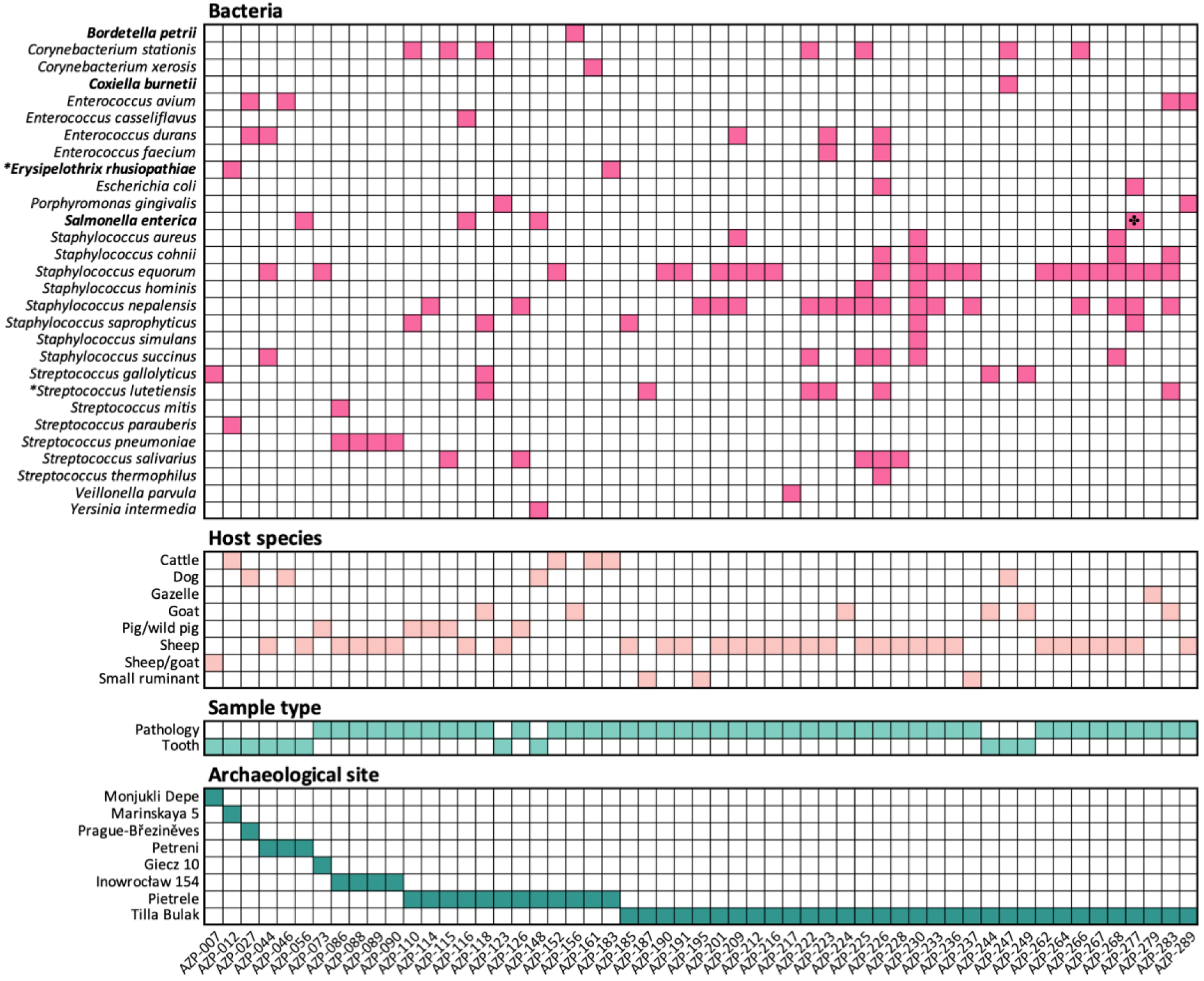
Host, sample type, and archaeological site information for all identified bacterial pathogens. Strictly pathogenic bacterial species in humans shown in bold. * Marks species with sufficient data for phylogenetic placement. ✤ Denotes the sample, which passed our stringent quality criteria at the taxonomic id of *Salmonella enterica* subsp. *enterica* as well as the ancestral taxonomic node of *S. enterica*.

### Interpretation of DNA evidence in its palaeopathological context

All 55 samples that produced robust bacterial hits came from sites palaeopathologically investigated. Eleven of those samples were teeth (8.3% of all teeth sampled), while the remaining 44 were bones with palaeopathological lesions (23.3% of all bones with lesions). Considering only sites studied for palaeopathologies, this suggests that the targeted approach was more likely to yield DNA from pathogens (p-value<0.0002, Chi square test), a signal robust also when excluding teeth, which showed no palaeopathologies (p-value<0.006, Chi square test). The fact that no ancient pathogenic bacteria were authenticated from the 25 bones sampled without any detectable lesions (p-value<0.005, Chi square test) emphasizes the advantage of palaeopathological investigations for prioritisation of specimens. The majority of hits were to skeletal elements from sheep (n=31). Other host species included cattle, gazelle, goat, wild or domesticated pig, dog, and the more generalised ‘small ungulate’ category. The lesions which produced bacterial hits were located on multiple types of skeletal elements, but most frequently on mandibles (n=8), vertebrae (n=8), and ribs (n=7), which is consistent with the skeletal elements we most frequently collected during the palaeopathological selection.

The oral cavity is a reservoir of commensal bacteria that can turn into opportunistic pathogens, but it is also an entry point for foreign bacteria ^41^. The pathological changes in the mandibles that produced bacterial hits included periodontitis, abscesses, and antemortem tooth loss, with genomic signatures for *Bordetella, Corynebacterium, Enterococcus, Salmonella, Staphylococcus,* and *Streptococcus* (**Figure 3, Table S2**). Periodontitis, for example, is a polymicrobial disease that is often caused by members of the oral microbiome, known as the ‘red complex’, including *Porphyromonas gingivalis*, which is the principal cause of this condition in humans ^40^ and known from broken-mouth periodontitis in sheep ^42^. We recovered DNA from *P. gingivalis* from a tooth, underlying the history of periodontal disease among human-associated livestock.

Sampled postcranial skeletal elements were primarily ribs and vertebrae. Lesions on ribs could be indicative of pulmonary infections, and several of the bacterial species signatures (n=7) recovered from lesions on ribs were species of *Staphylococcus* and *Streptococcus*, including *Streptococcus pneumoniae*, which colonise the airways and can cause infections in the lungs ^43^. Other species signatures recovered from palaeopathological lesions on ribs included species of *Enterococcus* and *E. coli*. The eight vertebrae that produced bacterial hits displayed pathologies characteristic of inflammatory disease and infection, with genomic matches to *Corynebacterium*, *Staphylococcus* and *Streptococcus*. However, whether the palaeopathological lesions and the ancient bacterial DNA signatures can be directly linked is unknown.

In terms of geographic patterns, more than half of the specimens with identified bacterial hits originated from the Tilla Bulak site (2000-1800 BCE) in Uzbekistan (Central Asia). Although the specimens collected from this site constitute 29% of the entire dataset, the number of positive bacterial hits (58.2% of total samples with bacterial hits) is disproportionate compared to other sites (p-value<0.0001; Chi square test). No other site was enriched for samples with pathogen signatures. While, we observe no correlation between the percentages of host and pathogen DNA recovered (**Figure S1**) and host DNA preservation was overall low (median 0.07%; mean 0.942% of QC passed reads), we identify, again, in samples from Tilla Bulak (27/105) as well as the Polish site Inowrocław (9/17) an elevated host DNA preservation of above 1% (p-value=0.0001 and p-value<0.0001, respectively; Chi square test; **Table S2**). This suggests that DNA preservation may have been overall better at the Bronze Age sites Tilla Bulak and Inowrocław, especially compared to the samples from the much older Pietrele site (4550-4250 BCE), which were also studied for pathology, processed similarly in the laboratory, and composed 23% of the dataset and yielded 22.2% of the bacterial hits. While age, deposition, and animal husbandry differed across sites as well as sequencing depth, it cannot be excluded that there was a higher pathogenic pressure during the period of deposition at Tilla Bulak compared to the time periods and geographical locations of the other sampled sites.

In sum, despite being mostly unspecific about the precise bacterial species, palaeopathological lesions can guide the prioritisation of promising bone specimens for the genomic investigation of pathogens. Moreover, our results suggest geographic and presumably site-specific differences in bacterial DNA preservation exist, which may be driven by variation in ante-or post-mortem treatment of animal remains impacting DNA preservation.

### Phylogenetic placement of zooarchaeological pathogens

To understand the evolutionary relationship of the identified bacteria and further confirm their ancient authenticity, we aimed to estimate their phylogenetic placement. While the low coverage impeded a complete genome reconstruction, we instead placed ancient genomes with large amounts of reads into the genetic diversity of modern representatives of that same species. Therefore, we ascertained mutations among contemporary genomes of the bacterial species together with an outgroup. For each position identified as variable among the contemporary genomes we set the base call for the ancient genome when all reads support a single base call (see Methods). While this approach does not allow us to infer the molecular evolution exclusive to the ancient genomes, it is informative about its phylogenetic placement to estimate the branching pattern and evolutionary relatedness to the modern variability ^31^.

Here we explore the phylogenetic placement for two identified species known to be pathogenic among livestock and for which we observed sufficient reads: *Erysipelothrix rhusiopathiae* and *Streptococcus lutetiensis*. First, *E. rhusiopathiae* is a multihost zoonotic pathogen with a significant economic impact particularly on swine as well as cattle production, but also as an occupational pathogen in humans ^38^. We recovered an authentic ancient DNA signature in AZP-012 with over 19,000 reads (0.5 coverage, 38% breadth) being aligned to the reference genome 52683_D03 (**Figure S2**, **Table S5**). AZP-012 was sampled from a cattle tooth excavated at Malinskaya 5 (Russia) dated to 2100 - 1800 BCE. Ascertaining 11,970 mutations among the outgroup *E. tonsillarum*, a previously human-derived medieval *E. rhusiopathiae* genome and 42 *E. rhusiopathiae* genomes isolated from a diverse set of contemporary hosts (**Table S4**), including domesticates and marine mammals, we infer the ancient genome is phylogenetically placed at the base of a sub-clade within *E. rhusiopathiae* independently defined by three recently published ancient genomes from medieval human remains (**Figure 5, Figure S3**) ^37^. The bootstrap value for the node separating the ancient samples from the closest modern genomes is only 67, driven by the resampling process and the relatively few mutations (23) identified among all ancient samples separating the ancient clade from the modern samples. However, the Transfer-Bootstrap-Expectation ^44^ value of this node is high (95.7, **Figure S4**), indicating that the overall placement of the ancient genomes is highly consistent, even though the exact reconstruction of the ancient sample placement is unstable. Nevertheless, the phylogenetic placement of AZP-012 together with independently generated ancient genomes from Iberian human remains suggests *E. rhusiopathiae* was a widespread multi-host pathogen in the past, harboring within-species diversity not yet described among modern representatives.

**Figure 5:**
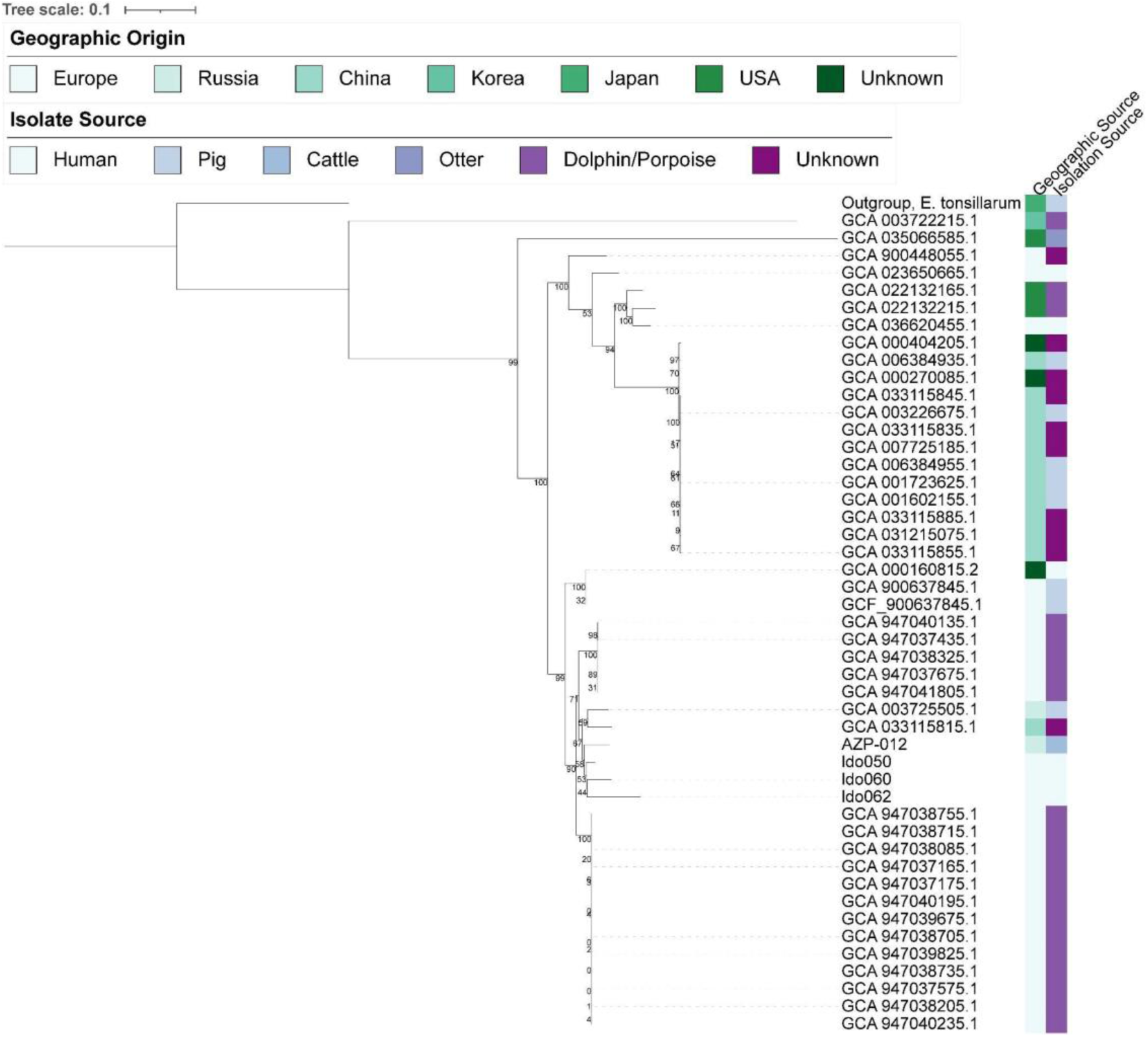
Phylogenetic placement of AZP-12 within the *E. rhusiopathiae* diversity. Among 42 *E. rhusiopathiae* genomes and a single *E. tonsillarum* (outgroup) genome we ascertained 11,970 mutations. For four ancient genomes (AZP-012 and three previously published human-derived genomes) the nucleotide call was inferred for each of the mutations (**Table S5**), to place the ancient *E. rhusiopathiae* sequences into the phylogeny. AZP-012 from Marinskaya 5 (North Caucasus) relates basal to the other ancient samples, as expected based on the age of the samples. Felsenstein Bootstrap support (of 100 replicates) for each node is displayed. Transfer-Bootstrap-Expectation values are shown in **Figure S4**.

*S. lutetiensis* is a member of the *Streptococcus bovis* type subgroup, along with the species *S. equinus*, *S. gallolyticus*, *S. infantarius*, and *S. salivarius* as an outgroup ^45^. *S. lutetiensis* has been primarily described to cause bovine mastitis, a costly disease impacting dairy cattle worldwide, but has also been isolated from other domesticates and humans ^46,47^. After alignment to the reference genome 45473_D02 we identified authentic *S. lutetiensis* signatures in three samples: sheep AZP-223 (0.18 coverage, 6.4% breadth), sheep AZP-226 (1.5 coverage, 70% breadth), and goat AZP-283 (0.11 coverage, 7% breadth), all excavated at Tilla Bulak (Uzbekistan) dated to 2000 - 1600 BCE (**Figure S2**).

In order to place the ancient genomic information into the modern genetic diversity, we leveraged genomes of *S. lutetiensis*, *S. infantarius*, and *S. gallolyticus* isolated from different sources across Eurasia (**Table S4**). We identified 92,526 phylogenetically informative positions across the modern representatives. All three ancient genomes are placed monophyletic, basal to the *S. lutetiensis* diversity (**Figure 6**, **Table S5**). Again, the low number of known mutations, 23, identified across all ancient samples separating the three ancient genomes from the modern diversity leads to instability measured by the classic Felsenstein bootstrap (97 - 47); however, their overall placement basal to modern *S. lutetiensis* is stable using the Transfer-Bootstrap-Expectation (97, **Figure S5**) and single-ancient-genome projections are placed consistently in 100% of Felsenstein bootstrap realizations (**Figure S6**).

**Figure 6:**
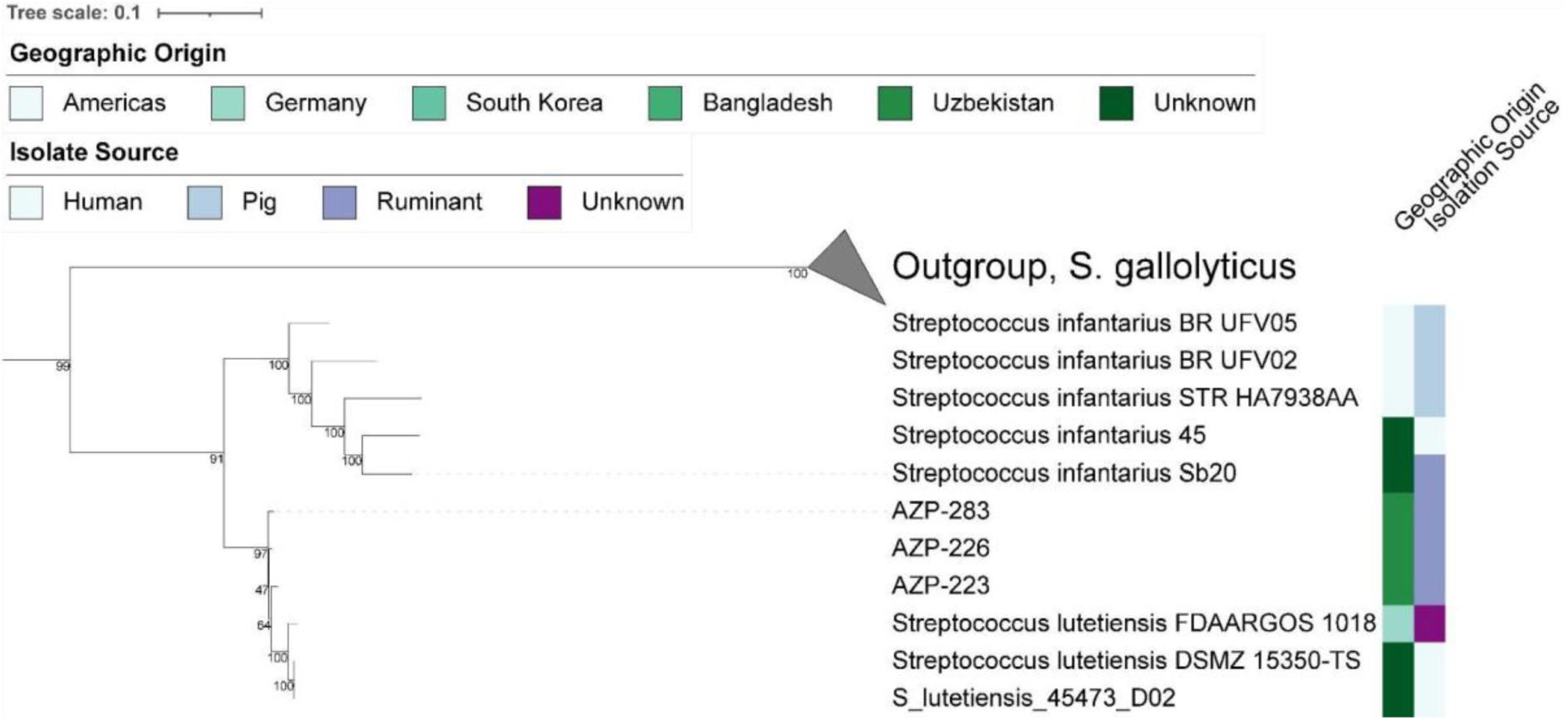
Phylogenetic placement of AZP-283, AZP-226, and AZP-223 within the *Streptococcus* bovis-type subgroup. The diversity (92,526 mutations) was ascertained using in total 21 *S. lutetiensis, S. infantarius, and S. gallolyticus* reference genomes mapped to *S. lutetiensis* 45473_D02 (**Table S4, Table S5**). All three ancient genomes originate from the Central Asian site of Tilla Bulak and form a monophyletic group basal to the known diversity of *S. lutetiensis*. Felsenstein bootstrap values are shown, the Transfer-Bootstrap-Expectation is presented in **Figure S5**.

Taken together, the basal phylogenetic placement of *S. lutetiensis* and the clustering of the ancient genomes from *E. rhusiopathiae* further corroborates the ancient authenticity of the identified pathogen DNA and provides first evidence for the widespread distribution and deep evolutionary history of both pathogens with domesticated animals. Nevertheless, the analyses of these pathogens are restricted to a few host species due to the limited representation in available databases, which prevented the phylogenetic projection of other candidates, like *Staphylococcus nepalensis*. As such, our inference is only a first step into the largely uncharted territory of prehistoric infectious disease in animals and livestock, and calls for further investigation of zoonotic pathogens both through the zooarchaeological record and with increased surveillance in modern contexts.

## Discussion

Prehistoric disease reservoirs in faunal populations and the likely increase in pathogen exposure that followed animal domestication are poorly understood. In this study, we applied a targeted approach to explore the suitability of zooarchaeological remains that primarily displayed palaeopathological lesions, which could have resulted from infection, for genomic palaeomicrobiological analysis.

Our results suggest that leveraging samples with palaeopathological lesions for genomic analysis provides a focused and valuable method for identifying ancient pathogens from zooarchaeological remains. Here, it is important to recognize that animal palaeopathology differs from human palaeopathology, both in terms of material studied, but also in the conclusions that can be drawn from morphological analysis alone. A key challenge is the disarticulated and fragmented nature of most zooarchaeological deposits, which precludes the systematic, skeleton-wide palaeopathological analyses typically performed on human remains ^20,48^. Moreover, skeletal pathologies in animals in the past have historically been understudied, with limited support from modern data, leading to continued reliance on assumptions derived from human pathological conditions ^20,48^. As a result, palaeopathological analyses of animal remains are therefore largely limited to isolated skeletal elements, which restricts analyses to the characterisation of a general health status. All of these complicate the formulation of a differential diagnosis, thus preventing the identification and study of specific diseases within and across populations. Compounding this issue, few zoonotic infections leave diagnostic traces in animal bone assemblages, making infectious disease in animal populations difficult to identify through morphological analysis alone ^49^.

While we recovered ancient pathogen DNA from both teeth and bones with palaeopathological lesions, our results suggest that sampling lesions offers advantages over random sampling in pathogen DNA retrieval from zooarchaeological remains. Nevertheless, targeting palaeopathological lesions for pathogen DNA retrieval has several limitations. Lesions are areas of bone remodelling and degradation, which is less optimal for DNA preservation than dense bone in agreement with the overall poor host DNA preservation. Lesions are often more exposed to environmental factors resulting in post-mortem fractures, or intrusions of soil that can be difficult to remove, while pretreatment methods like bleach or UV radiation risk destroying pathogen DNA. Additionally, the lesions themselves may not be the optimal sampling locations for certain pathogens, but the absence of a differential diagnosis inhibits the customisation of a DNA sampling strategy based on the pathophysiology of the pathogen that caused it. Moreover, we observe variation in DNA preservation across sites presumably due to differences in depositional practices, environmental conditions, and specimen handling, all of which may contribute to the differences observed in pathogen DNA recovery. Future investigations explicitly designed with comparable amounts of pathological and non-pathological specimens per site will be instrumental to further corroborate our findings.

Bone remodelling is generally a slow process ^50^; although, some acute infections can cause skeletal changes, i.e., acute osteomyelitis. As a result, palaeopathological lesions may reflect both long term (chronic), and less frequently, acute infections ^51–53^. While the identified bacteria cannot be directly linked with the broadly defined lesions, they may represent opportunistic colonization of an existing site of infection. Moreover, we do not recommend focusing exclusively on palaeopathology when designing a sampling strategy for ancient pathogen retrieval from zooarchaeological remains as doing so risks excluding the many pathogens responsible for acute infections. Individuals, who perished from acute infections, are often not visible in the archaeological record, as these infections do not leave marks on the bones ^54,55^, but many ancient pathogens from acute septic infections have been recovered from the dental pulp chambers of humans without any osteological evidence of disease^21,22^.

We observe this phenomenon in our dataset by the detection of *S. enterica* signatures, a group of bacteria responsible for gastroenteritis or systemic disease in humans and animals ^32,33,56^, in four specimens from three sites across Eurasia covering a period between 4,000 to 8,000 years ago. Interestingly, two of the four signatures are from teeth, one is from a lesional mandible, and one is from an arthropathic lesion on a femur, pointing at additional skeletal material that may serve as a source for ancient *S. enterica* DNA. The emergence of human salmonellosis has previously been linked to the Neolithization of Eurasia and likely has a zoonotic origin ^22^. Although the screening dataset did not provide sufficient genome coverage for further analysis of this bacterium, future deep sequencing or target enrichment capture may provide more clarity on the relation between these animal-derived hits and known ancient and modern *S. enterica* diversity.

We also detected other known zoonotic pathogens, where *C. burnetii* ^35^ and *E. rhusiopathiae* ^36^ are widespread occupational pathogens today. The identification of several human pathogens from the zooarchaeological record highlight the potential for zoonotic disease emergence as a risk factor during the Neolithization across Eurasia. Consistent with this, Sikora et al. ^25^ identified an uptick in zoonotic pathogens in particular among pastoralist communities associated with animal husbandry, underlying the drawbacks of our intimate relationship with domesticates. Although most of the authenticated bacterial genomes in this study had too low coverage for further analysis, the four samples used in the phylogenetic analyses demonstrate that the authentication process is robust. While such analyses were not possible for all potential pathogens detected here, they are important in order to distinguish between signatures of authentic endogenous pathogens and any, especially opportunistic pathogens present in the human microbiome, that may have been deposited during excavation and subsequent handling, and could have developed degradation patterns reminiscent of authentic ancient DNA while in storage.

Ancient DNA offers a powerful means of detecting infectious agents not only in pathological bone with non-specific lesions but also in skeletal elements absent of pathology or where pathology has been obscured by taphonomic processes. By enabling the recovery of pathogen DNA from these remains, ancient DNA analysis provides a more comprehensive picture of past zoonotic infections and their evolutionary trajectories, helping to uncover the long-term dynamics of host-pathogen interactions and the role of animals in disease transmission. Altogether, while targeting palaeopathological lesions can provide valuable insights, it is not sufficient for a comprehensive understanding of ancient infectious diseases. Hence, this study is a first step towards exploring ancient pathogens in the zooarchaeological record, which holds the promise to elucidate the host range of zoonotic pathogens and the genetic mechanisms enabling the evolutionary adaptation towards the human host.

## Materials & Methods

### Material sources, palaeopathological investigation and site selection

The study materials were animal bones and teeth. An initial archaeozoological assessment had been carried out for all the faunal assemblages. Although human (brachydont) teeth have been shown to preserve DNA from blood-borne pathogens ^31^, it is unclear if the same is true in species that do not have enclosed pulp chambers, such as ruminants (hypsodonts ^57^). Teeth (N=132) were collected from all sites except those in Poland and two in Germany. Bones were collected from Mikulovice (Czech Republic), the German sites, the Polish sites, Pietrele (Romania), and Tilla Bulak (Uzbekistan). Palaeopathological investigations were carried out only on the Polish, German, Pietrele, and Tilla Bulak sites (**Table S1**). The analysis of pathology was conducted using a wide range of references and classifications ^19,58^ lesions that displayed signs of an active infection ^19,58^. No selection was made based on wild or domesticated animals, but given the post Neolithic ages of both sites, the abundance of skeletal elements from domesticated species outweighed those from wild species. All selected bones from Poland, Pietrele, and Tilla Bulak and some of the bones from Germany displayed pathological lesions (N=189; **Table S2**).

The choice of sites was dictated by several factors. Their different chronologies made it possible to capture changes over time and, in particular, the timing of pathogen transmission in the human–animal relationship. The geographic range of the sites could help indicate the potential location of the emergence of zoonotic disease. In addition, each of the sites is set in a different environmental context, and this variety provided the opportunity to avoid biases due to the environment on DNA and palaeopathology preservation potentially present at single sites. All sites are listed in **Table S1** and a summary of the sites can be found in the **Supplemental Online Material**.

### Lab processing

#### Drilling

All samples were processed in dedicated clean room facilities. Aside from the German samples, which were processed entirely at the ancient DNA facility in Tartu, Estonia, all selected samples were drilled in the ancient DNA laboratory at the MPIIB in Berlin.

With the exception of the German samples, only one sample was collected from each element. For some of the German specimens multiple subsamples were collected, and when mandibles or maxillae included teeth, these were also sampled. Occasionally, multiple teeth were collected from the same individual and analysed as separate subsamples (**Table S2**).

The roots of all teeth selected for the study were sawed off, and powder was collected by drilling into the root, the pulp cavity, or both depending on the morphology and size of the tooth. Due to their size, small/ milk teeth were wiped with bleach solution, cleaned with water, and then pulverised entirely using a mortar and pestle. Pig teeth had very fragile roots that could not be drilled without fragmenting while sheep/ goat teeth lacked a pulp chamber.

Palaeopathological lesions were sampled directly. No pretreatment of bone was applied in order to prevent potential destruction of any pathogen DNA present in the lesions. For each lesion, drilling was carried out across the surface of the lesion and, where possible, this was complemented with an additional sampling by drilling deeper into the cancellous bone beneath the lesion. When possible, trabecular bone was also collected.

Bones without lesions were sampled in a similar manner to those with palaeopathological lesions. This included drilling across the surface and then deeper into the bone. If possible, a sample from trabecular bone was included.

Approximately 50 mg bone or tooth powder was collected from each sample. In cases where larger amounts of powder had been collected, samples were pulse vortexed three times and then 50 mg of the mixed powder was weighed out.

#### DNA extraction, libraries, and sequencing

DNA extraction was carried out using methods optimised for ancient DNA ^59,60^. 1 mL of lysis buffer consisting of 5M EDTA (pH 8.0), Tween 20, Proteinase K, and H_2_O was added to each 50 mg bone or tooth sample. The tubes were then sealed with parafilm and incubated with rotation overnight at 37°C.

Following lysis, samples were centrifuged for 2 Minutes at 16,400 g to pellet any undissolved material. The supernatant was collected and transferred to 10 mL of binding buffer (5 M guanidine hydrochloride, 40% (vol/vol) 2-propanol, 0.12 M sodium acetate, and 0.05% (vol/vol) Tween 20). Samples were then filtered through minElute columns and washed twice with 200 μL PE buffer (Qiagen) before elution in 50 μL EBT. DNA concentration of the extracts was determined using the Qubit HS DNA kit.

Libraries constructed in Tartu followed the Meyer and Kircher protocol for double-stranded DNA ^61^. In Berlin, single-strand DNA libraries were constructed following the Santa Cruz Reaction protocol ^62^. A qPCR test was performed to determine the optimal number of PCR cycles for each sample using Maxima SYBR Green/ROX qPCR Master Mix (Thermo Scientific) following the set-up recommended by the manufacturer on a OneStepPlus machine (Applied Biosystems). Dual indexing PCR was performed using AmpliTaq Gold 360 Mastermix (Applied Biosystems). PCR products were purified using a 1.2 ratio of Sera-mag speed beads (Fisher Scientific). Beads were washed twice with 200 μL ethanol (80%) and eluted in 30 μL EBT. Final DNA concentrations were measured using the Qubit 1X dsDNA High Sensitivity kit (Invitrogen), and average fragment lengths were measured on a Fragment Analyzer (Agilent) using the High Sensitivity NGS Fragment kit (Agilent). Samples were pooled based on equimolarity and sequenced on Novaseq SP flowcells in 50PE mode (Berlin) or the Illumina NextSeq 500 platform in 150PE mode (Tartu). Extracts and libraries can be made available upon request for further investigations.

### Bioinformatic processing

#### Screening pipeline

nf-core/eager was used for bioinformatic processing, including quality control, initial mapping to the human reference genome, and finally metagenomic screening of unmapped reads for candidate pathogens using MALT and HOPS ^28,29^. See Raw Data Processing in project github for parameters. A list of taxa of candidate pathogen species and genera can be found in (**Table S3**), which includes known zoonotic pathogens and genera, along with additional animal-specific pathogens. We removed the genus node of *Bacillus*, *Clostridium* and *Brucella* due to closely related non-pathogenic species highly abundant in soil that complicates the identification of positive samples. A custom screening database based upon the RefSeq ^63^ was used as described previously ^14^. A custom HOPS-post-processing script was used to identify candidate samples (Supplemental Information), which allows for the manual setting of various heuristic thresholds and output was limited to samples with pathogen hits that fulfil all of our set criteria. The criteria were: (1) a minimum number of 50 reads assigned to a taxonomic node of interest, (2) a default edit distance ratio of 0.8, (3) an ancient edit distance ratio of 0.5, and (4) a minimum read distribution of 0.6, and (5) damage on 10% of reads with possible damage transition patterns (C→T) at the 5’ end; or, also the 3’ end (G→A), in the case of double stranded libraries. The read distribution metric (number of covered genome positions / sum of aligned read lengths) ^28^ is designed to be highly attuned to detecting particularly weak pathogen signatures samples, since the value decreases with any read stacking which would be expected from spurious misalignments by reads from organisms with homologous genome segments within the metagenomic sequencing library, a major concern for false positives. However, this metric is inappropriate for high values of reads assigned to a specific candidate pathogen node, since with many reads, we expect some measure of overlap, leading to a reduced read distribution value. To ensure that we identified any pathogen signatures constituted from many reads for which the read distribution cutoff would be inappropriate, we separately ran the post-processing script with the same requirements, except for minimum number of reads >10,000 and no read distribution cutoff.

#### Host validation with kraken2

The taxonomic profiling pipeline nf-core/taxprofiler was used to confirm, identify, or correct the host identity of each metagenomic dataset ^64^. We performed short-read sequencing quality control (--perform_shortread_qc) and utilised kraken2 with a prebuilt nt database dated 11/29/2023 retrieved from https://benlangmead.github.io/aws-indexes/k2 to taxonomically assign all reads that passed QC using the--kraken2_save_minimzers option. Metagenomic host classification can be challenging due to high variability in sample preservation and contamination with environmental DNA, both of which interfere with reliable taxonomic host identification. Moreover, several eukaryotic species are regularly identified in ancient metagenomic datasets, for example European hedgehog (*Erinaceus europaeus*), suggesting contamination also at reference genome level leading to spurious results ^65^. To identify the host genus, we identified the top genus taxonomic nodes below the Metazoa kingdom node and set two cutoffs for assessing the host identity using metagenomic taxonomic classification. First, we set a minimum threshold of 0.1% of input QC-passed reads assigned to a given taxonomic node for it to be trusted as a host identification when the host genus was in agreement with the prior morphological identification, when available. A total of 39.4% of samples reached that threshold of which 77.9% agreed with the morphological host identification. Values below 0.1% of a single genus assignment indicate poor sample preservation at the endogenous DNA level and were not considered further for taxonomic identification. Second. a threshold of 1% of reads was required to update a host identity in the case that the morphological host identity was inconsistent with the metagenomically-assigned top genus, unless the metagenomically-assigned top genus was *Homo*, which presumably stems from contamination. Only 14.6% of samples reached that threshold of which 17.3% of the samples host species was updated. When no morphological host identification was possible, or multiple potential hosts were proposed, the 0.1% threshold was used for metagenomic host genus assignment. Results are shown in **Table S2**. No correlation between host DNA and pathogen DNA preservation was observed (**Figure S1**).

#### Pathogen phylogenetic reconstruction

In order to validate our metagenomic microbial screening results and learn about the evolutionary history of the identified ancient pathogens, we performed phylogenetic projection of the ancient genomes into the known modern diversity of the respective species. We focused our effort on two species - *E. rhusiopathiae* and *S. lutetiensis,-* with at least one sample with a high number of reads (3402 - 7476) assigned specifically to the species node during metagenomic screening, which present the greatest likelihood of having enough genetic data for successful phylogenetic projection, had known pathogenic impacts, and whose species diversity was described.

For each of the two species, we utilised a set of modern genomes to build a mutation table representing the known genetic variation of that species. Selected reference data and associated metadata is available in **Table S**4. For samples from GenBank without any available raw sequencing reads, we utilised wgsim (https://github.com/lh3/wgsim) to simulate sequencing data to map back to the reference genome. Additionally, for *E. rhusiopathiae*, we reanalyzed publicly available ancient metagenomes previously identified as positive for the species in Iberia from the past 1000 years ^37^ with the ancient metagenome we recovered (AZP-012). Initial data processing including adapter trimming, low quality read filtering, mapping and deduplication was carried out by nf-core/eager separately for the modern and ancient samples (See Raw Data Processing in project github for parameters) for each species to allow for ancient DNA specific mapping parameters and subsequent masking of terminal bases most likely to be affected by ancient DNA damage to be used for the metagenomic data. Deduplicated bam files were used as input for an in-house Snakemake pipeline which collects all variable positions along with support statistics across all samples for downstream processing using an interactive python environment ^66^.

For each species, we performed stringent quality control to obtain a table containing the mutations present among the modern genomes. First, we removed any modern genome with an average coverage below 4. Second, the nucleotide call in any modern sample was filtered and set to N if it failed any of the following criteria: the FQ score lower than-30, the major allele frequency of the position lower than 90%, fewer than 2 forward reads, fewer than 2 reverse reads, fewer than 4 reads total, or if more than 50% of reads supporting a base call also supported an indel call within 3 bases. Lastly, we subset the positions for downstream analysis to only the core genome, which we defined as positions called in 100% of modern samples analysed. Then, we built a single nucleotide polymorphism (SNP) table of phylogenetically informative positions which varied in at least one modern sample or ancient sample with >1x coverage. Importantly, for all ancient genomes with <1x coverage (all but ldo050 for *E. rhusiopathiae*), we exclude identifying phylogenetically informative SNPs from ancient samples (singletons), which could be biased away from the modern genomes due to metagenomic mismappings and are hard to call correctly. *E. rhusiopathiae* (sample AZP-012) or *S. lutetiensis* (sample AZP-223, AZP-226, AZP-283), attained a genome-wide coverage of >0.05x and hence were projected into the SNP-table generated using the respective set of modern genomes per species. For each retained variable site identified among the modern genomes, the ancient genotype was called in each sample with a coverage of ≥1, FQ score =-30 and major allele frequency >0.9. The ancient genotype was set to “N” for any site variable among the modern genomes which failed any of these criteria. The complete SNP tables are available on the project github page (see Supplementary Information) and were used to generate a maximum likelihood (ML) tree using RAxML ^67^ with 100 bootstrap replicates.

To validate the phylogenetic placement of the ancient samples, we undertook two additional analyses. First, we calculated the Transfer-Bootstrap-Expectation metric using RAxML using 100 bootstrap replicates. This metric is useful for assessing the overall stability of a node relative to the rest of the phylogenetic tree. Second, to ensure that the high level of missingness in ancient samples (range 23-90%) was not biasing their phylogenetic placement, we created a ML tree with 100 bootstrap replicates separately for each projected ancient sample, with only sites called across all modern ingroup samples and the ancient sample included in the projection and variability within either the ingroup or outgroup samples. The topology of the ML realisation trees broadly recaptures the tree topology observed when allowing for missing data, and are consistent across different ancient genomes (**Figure S3** and **S6**).

Second, phylogenetic trees were visualised using iTOL and were annotated for isolation source and geographic source using publicly available metadata.

#### Statistical tests

Chi square tests were used for comparisons of preservation by site and pathogen recovery by site by comparing the number of samples from that site with or without >1% host DNA assigned or a pathogen identification compared to all other sites collapsed, respectively. Multiple testing correction was done by the Bonferroni method (adjusted alpha=0.00189). Only p-values significant after multiple testing corrections have been reported. Comparisons for palaeopathological lesions for given sites and for recovery of pathogen hits were conducted only on samples from sites that were assessed for palaeopathologies (**Table S1**).

## Data availability

DNA sequencing data is available at the European Nucleotide Archive under project accession PRJEB63473. Individual accession identifiers are also listed in **Table S2**.

Code and necessary files for recreating downstream analysis can be found at the project github repository (https://github.com/fm-key-lab/Zooscreen).

## Supporting information

Supplementary Figures 1-6, Archeological site descriptions

## Acknowledgements

FMK is supported by the Klaus Tschira Foundation (GSO/KT030) and the EASI-Genomics TNA project PID10336. FMK, AKWR, and ILM are supported by the Max Planck Society. The taphonomic and palaeopathological work (KP), was carried out as a part of two research internships at the Max Planck Institute (2022 and 2023: ID-UB Mobility program, Call No. 72 Task 7). We thank Rainer Linke (Königsbrunn) and Sebastian Gairhos (Stadtarchäologie Augsburg) for providing access to samples. Excavations at Tilla Bulak were funded by a grant from the Gerda Henkel Foundation, Düsseldorf, Germany (Az. 16/ZA/07). We would like to thank the German Archaeological Institute for providing work space for KP and AKWR. We would also like to thank Meike Sorensen for her help with testing the SCR protocol before it was implemented in the aDNA lab, and for all her support in setting up the lab. We are grateful to Diane Schad for her help with refining the figures. We thank the Key Lab for helpful discussion. Finally, we thank all the people who have been involved in the archaeological excavations from which we obtained samples, as well as those responsible for the curation of each site.

## Author contributions

FMK conceived the project with conceptual input from KP. KM, ST, JE, ME, RK, MH, DP, AC, NB, DD, JK, AN, AAK, ARK, VEM, ABB, MT, SH, PWS, KK, RU, SR, REG and KP curated the archaeological materials and provided contextual information. ST, MH, NB and REG performed initial zooarchaeological analyses. KP performed palaeopathological analyses (German, Polish, Pietrele and Tilla Bulak samples). EAN performed palaeopathological assessments of the German collection and designed sampling strategies for ancient DNA analysis. CK, KB shared expertise in palaeopathology. AKWR, MK, HK and CS performed the ancient DNA laboratory work. ILM analysed the data with input from FMK. AKWR, ILM, KP and FMK interpreted the DNA and palaeopathological results. All authors revised and accepted the manuscript prior to publication.

