## Supplementary Figures 1-6, Archeological site descriptions for "Probing the zooarchaeological record across time and space for ancient pathogens"

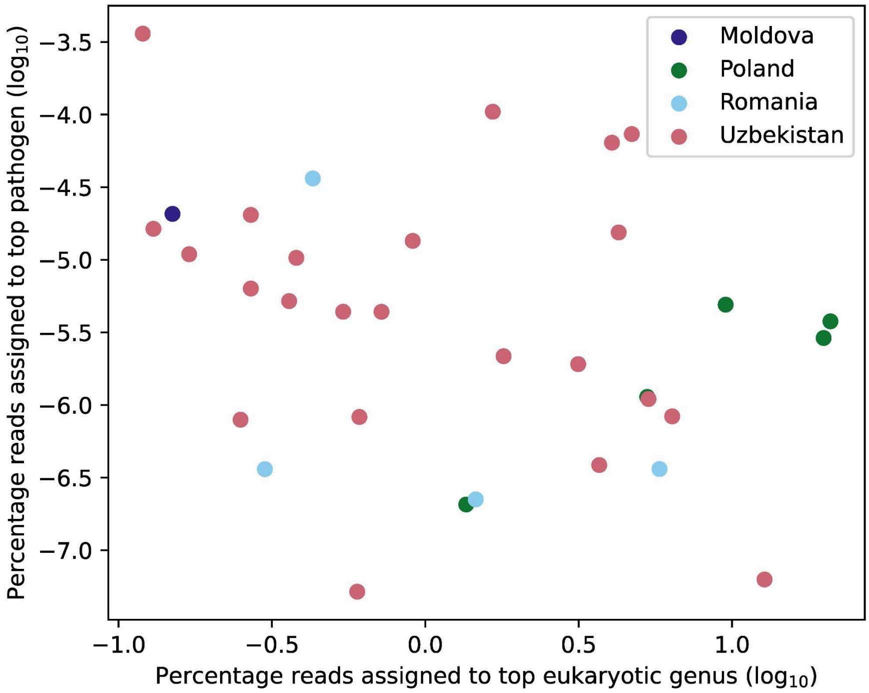


**Figure S1.** Percentages of reads assigned to top eukaryotic genus vs percentage of reads assigned to top pathogen species for samples with a pathogen identified. The pathogen with the highest number of reads was plotted for samples with multiple identified pathogens. No correlation is observed between pathogen and eukaryotic read percentages, suggesting that preservation of host DNA does not predict pathogen recovery ($R^2$=0.023)

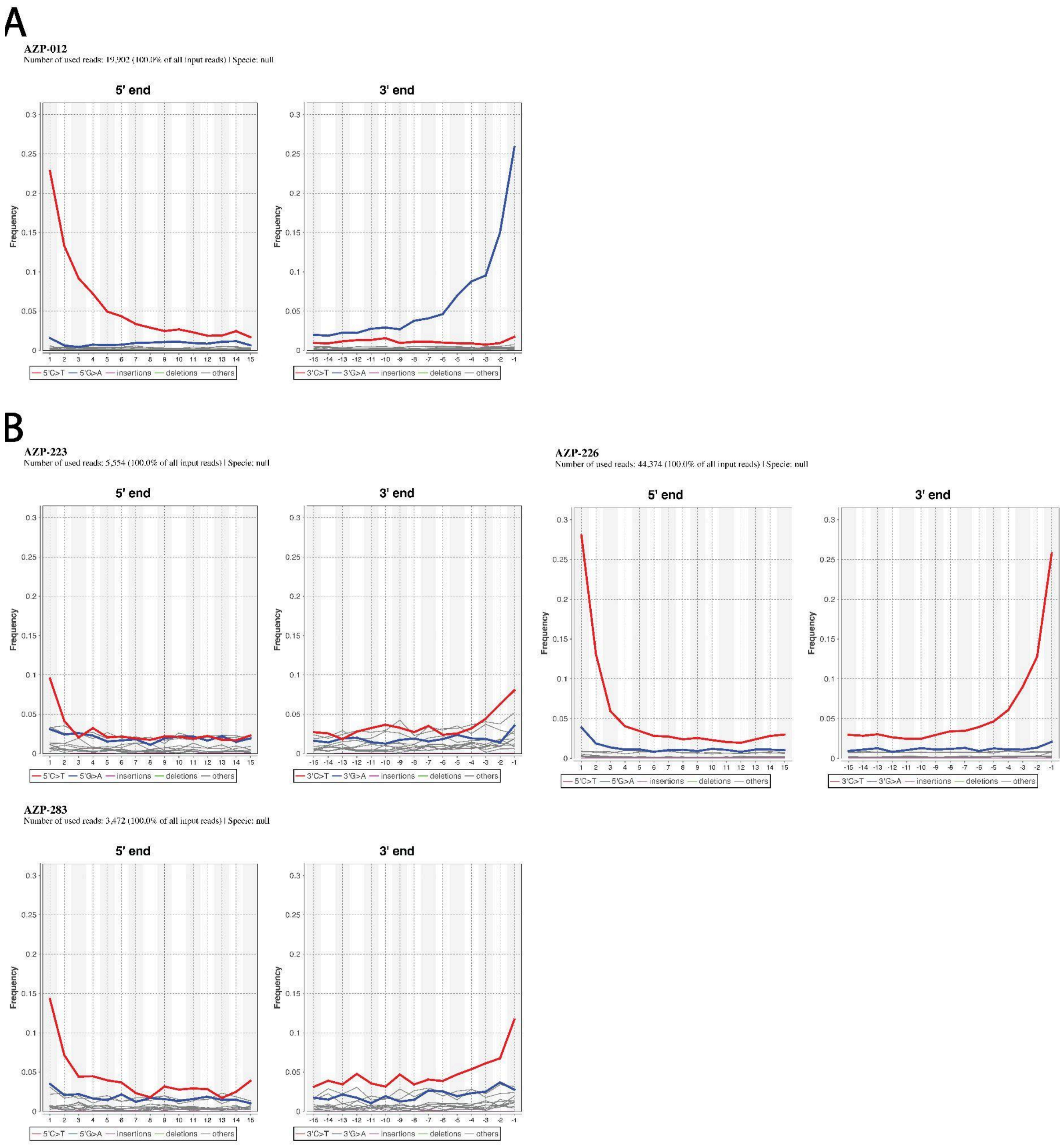


**Figure S2**. Damageprofiler plots display rates of base substitutions reflective ancient origin for A) *E. rhusiopathiae* and B) *S. lutetiensis*. AZP-012 was prepared using a double-stranded protocol and thus has C->T transitions on the 5' end and G->A transitions on the 3' end of reads. AZP-223, AZP-226, and AZP-283 display C->T transitions on both the 5' and 3' ends of the mapped reads, consistent with single-stranded library preparation.

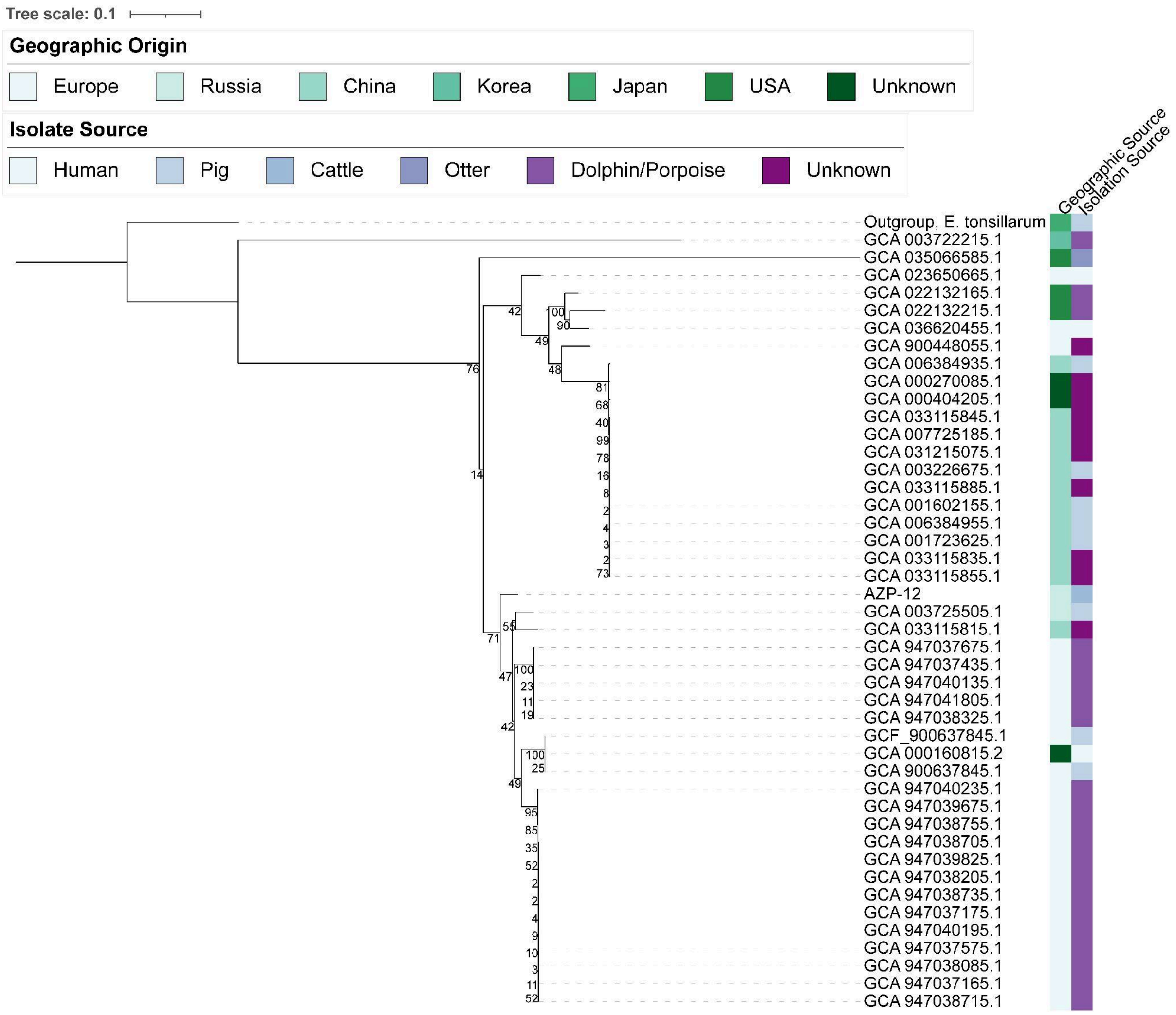


**Figure S3**. *E. rhusiopathiae* AZP-012 single sample phylogenetic placement. Previously published ancient genomes are excluded.

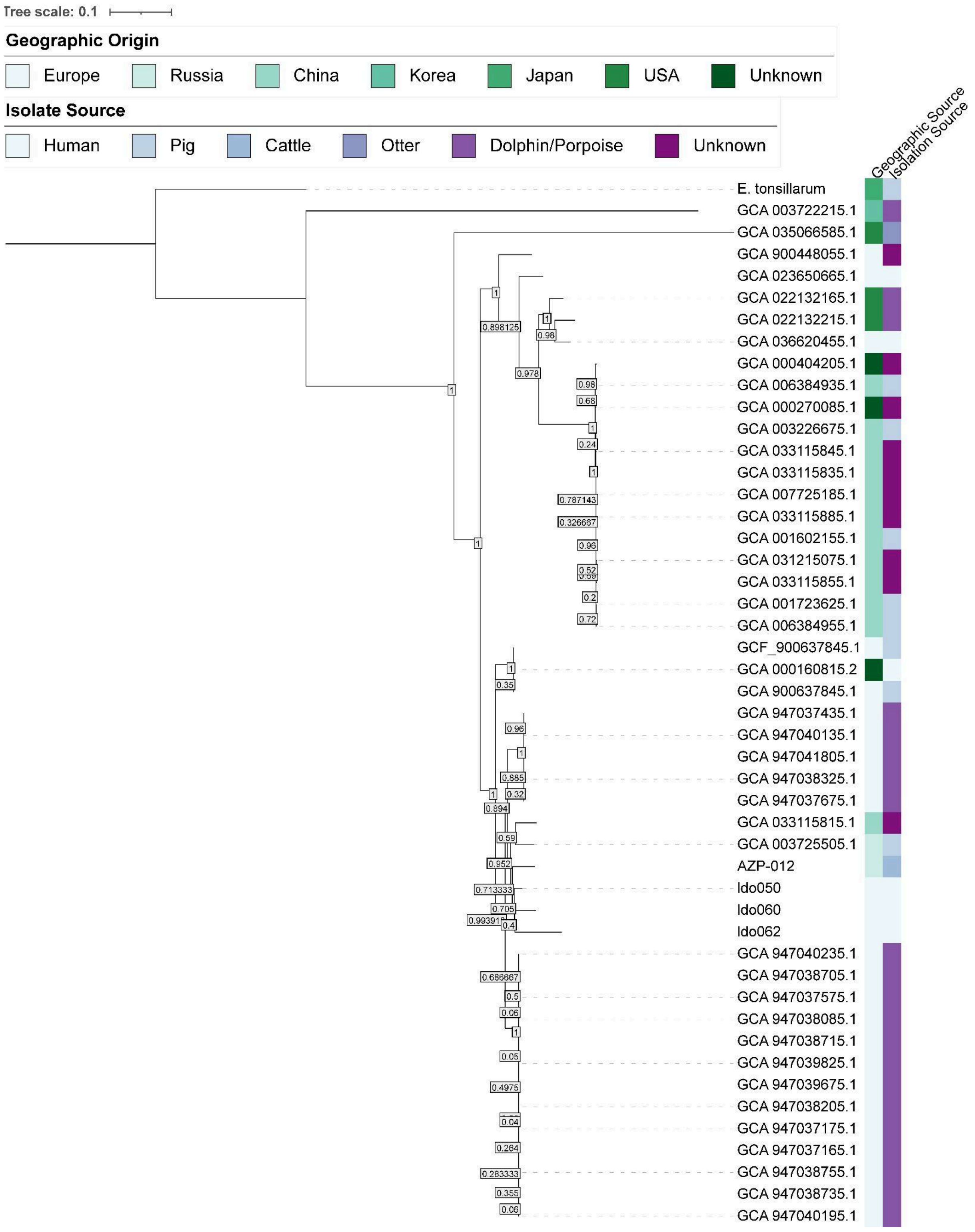


**Figure S4.** *E. rhusiopathiae* Transfer-Bootstrap-Expectation statistics. Bootstrap value of node separating ancient samples is 0.952, highlighted in bold.

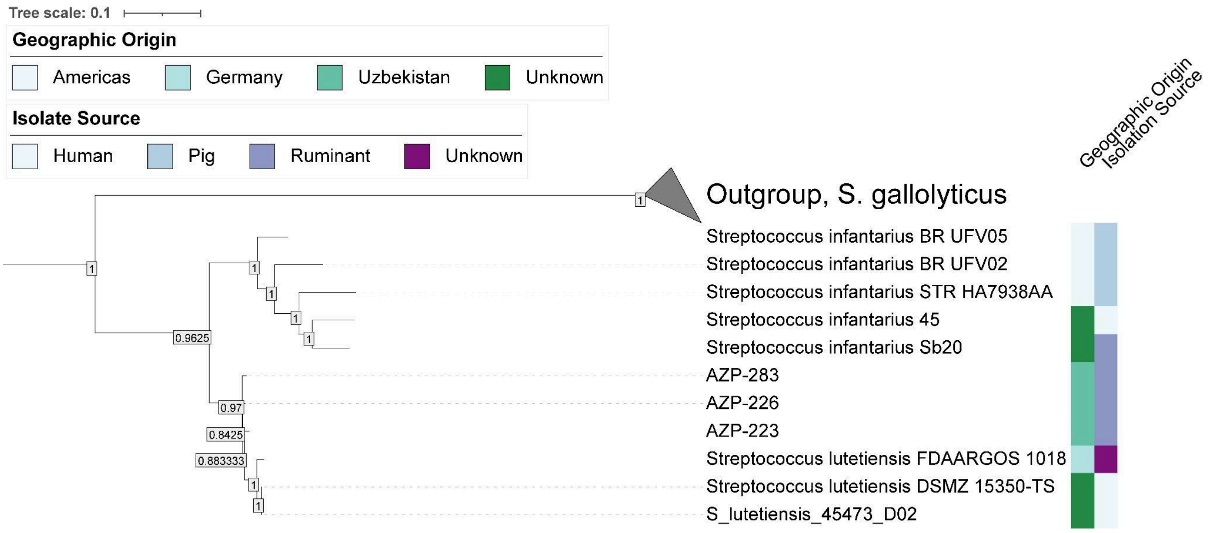


**Figure S5.** *S. lutetiensis* Transfer-Bootstrap-Expectation statistics. Bootstrap value of node separating ancient samples is 0.952, highlighted in bold.

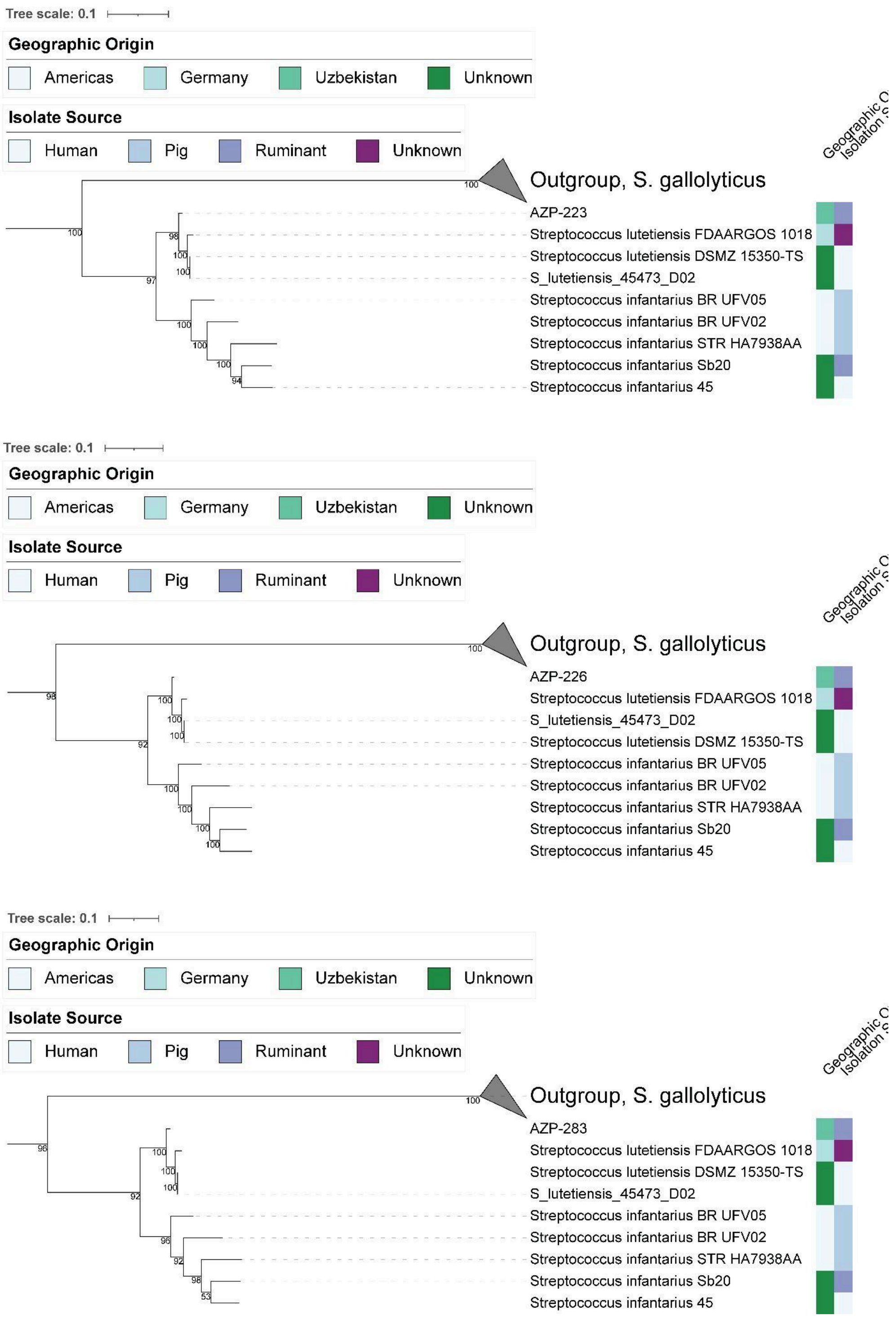


**Figure S6.** *S. lutetiensis* single sample phylogenetic placements.

### Materials and methods

#### Site, fauna, and pathology results

**Ken Massy, Simon Trixl, Jana Eger, Michal Ernée, René Kyselý, Michael Hochmuth, Dominik Poradowski, Aleksander Chrószcz, Norbert Benecke, David Daněček, Jana Klementová, Anatoli Nagler, Alexey A. Kalmykov, Anatoly R. Kantorovich, Vladimir E. Maslov, Andrey B. Belinskiy, Meda Toderaş, Svend Hansen, Philipp W. Stockhammer, Kai Kaniuth, Regina Uhl, Sabine Reinhold, Rosalind E. Gillis, Kamilla Pawlowska**

##### Czech Republic

###### *Dolní Beřkovice, Czech Republic*

The polycultural site at the merge of cadasters of Dolní Beřkovice and Křivenice (location Dýhárna Denzer) was excavated between 2001 and 2005 by the Institute of Archaeology of the Czech Academy of Sciences in Prague. It yielded a large number of animal bone remains (unpublished). The dog tooth analyzed part of this study is from a Corded Ware Culture (CWC) grave [1,2]. The human skeleton found in the grave 932 lay on the left side, which is a typical position for females. The sampled dog tooth (*caninus inferior sinister*) was excavated from grave 932 (ID: 5933) alongside a human skeleton laying on the left side, a position typical for females.

###### *Hostěnice (site Brozany), Czech Republic*

The site Brozany (Hostěnice) is located in lowlands in the marge of Central and North Bohemia and was excavated 1997 [3]. Osteological material [4] dated to Funnelbeaker Culture represents standard settlement waste but puppy skeletons and fox skeletons were also found. One dog (ID: 726) tooth (*premolar 3 inferior sinister)* was included for DNA analysis in this study. The tooth, likely an offering, was found in settlement storage pit No. 181A.

###### Hostivice-Palouky, Prague-West district, central Bohemia, Czech Republic

Contact persons: David Daněček, Jana Klementová

Rescue excavation was carried out by J. Klementová and D. Daněček (Museum Roztoky) in 2007–2008, 2010 to 2013 [5,6]. The excavated area covers 10 ha, more than 1,300 settlement pit features and a similar number of post holes have been found (2626 features together). The site was occupied during the Linear and Stroked Pottery cultures, the Funnel Beaker and Řivnáč cultures, Hallstatt (Ha C-D1), Roman Iron Age, Early Middle Ages. More than 330 graves were also uncovered, and burials/skeletons were also identified in sunken settlement features from the Funnel Beaker (Baalberge stage), Corded Ware, Bell Beaker and Knovíz cultures (B D – Ha A2), as well as from the La Tène period (Lt B1-C), Migration period, and Early to Late Middle Ages. A small group of graves, few other single graves and two settlement pits (1356/2008 and 1415/2008) as far as group of few other sunken features (1723/2011) interpreted as “ceremonial structure” and possible “foundation deposit” [7,8] are dated to the Bell Beaker Culture. Multiple human skeletons were sampled for ancient DNA and presented previously [9] and several human specimens positive for *Yersinia pestis* from grave 17/22 part of Hostivice-Palouky have been described [10]. Anthropological research was performed by M. Dobisíková (National Museum Prague).

##### *Libkovice, Czech Republic*

Archaeological research carried out in 2018 by the Institute of Archaeological Monument Preservation of Northwest Bohemia in Most (V. Sušická) focused on the southeastern part of the cadastre of the extinct village of Libkovice, Most district. The site is located in lowland near the former confluence of the Lomský and Radčický streams. In an irregularly oval grave pit (grave 233/18) the skeleton of an individual lying on the left side was found, identified archaeologically as a woman. An amber necklace containing at least 20 corals of various shapes and sizes was found around the neck of the deceased. Another necklace of several dozen large drilled animal teeth was hung in the rib cage area. In front of the body there was a belt consisting of at least fifteen rows of several hundred drilled animal teeth of various types and sizes [2]. One of the drilled dog teeth (*caninus inferior dexter*) was chosen as material for ancient DNA analysis in this study.

##### *Mikulovice, Pardubice district, east Bohemia, Czech Republic*

Contact person: Michal Ernée

The site is located in Pardubice district, east Bohemia, Czech Republic and was excavated during rescue excavation in 2006–2012 (J. Frolík, R. Sedláček). The site contains thousands of sunken settlement features, about 100 Early Bronze Age (EBA) graves, and three settlement features containing 109 skeletons were documented in several groupings. With exception of two Proto-Únětice graves (No. 95 and 96), the graves mostly belong to the Classic and Post-Classic stages of the Únětice culture. The inhumation burials are very rich in so-called “exotics”, especially amber artefacts, which are present in 28 graves. The EBA cemetery has been completely analysed and published as a monograph [11] that is available open access on AcademiaEdu https://www.academia.edu/43585678/Mikulovice_Early_Bronze_Age_Cemetery_on_the_Amber_Road). It contains the complete anthropological (P. Stránská, E. Zazvonilová), paleopathological (L. Vargová, K. Vymazalová, L. Horáčková), demographical, anthropomorphic (epigenetics; P. Velemínský, J. Cvrček) and osteological (R. Kyselý) analyses [11].

The two samples included in this study came from grave 42 (feature 2040). This grave contained the skeleton of a child (6,5–7,5 years): right-sided crouched burial, head towards the south, east-faced. Grave goods: small bronze chisel behind the head, simple bronze armring on the left-hand wrist, three amber beads and one drilled dog teeth by the waist, animal bone (pig scapula [12]) in front of the knees. Palaeopathological findings: enamel hypoplasia, cribra orbitalia sin., by aDNA screening identified plague [10]. Archaeological dating: Classic Únětice culture. Radiocarbon dating: MAMS-30479 (3550±19) 1950–1779 BCE cal 2-sigma. [11]. Pandora No.: MIB054. NM Prague Inv. No.: P7A 43141.

Samples:

AZP-021, Grave 42, context 5031, Dog, Caninus superior sinister, drilled tooth (fang), offering,

AZP-022, Grave 42, context 5032, Pig, Scapula dextra, post mortem fragmented, offering

##### *Mochov, Czech Republic*

The excavation of site Mochov 3 (location „Na Nehvizdku“) was carried out in 1980 by the Museum of Čelákovice city (J. Špaček). The sunken feature no. 17 was dated archaeologically to Salzmű̈nde phase of Funnelbeaker Culture by M. Zápotocký (unpublished). This dating was later confirmed by radiocarbon analysis of the sample from the dog skeleton (4669 ± 24 BP, cal 3519 – 3369 BC (95%); analysis CRL-20_168: unpublished data). A dog skeleton found in the feature together with a child skeleton represents a non-standard finding undoubtedly with ritual connotation, despite the nature of deposition of the dog body is not clear from the terrain documentation. One tooth (*premolar 4 inferior dexter*) from the dog skeleton was chosen for DNA analysis.

##### *Prague-Březiněves, Czech Republic*

Several graves of Corded Ware Culture (CWC) were excavated in Prague-Březiněves in 2006 (location Na Boleslavce) by ARCHAIA Praha z.ú. (K. Svoboda). The excavation represents a typical CWC cemetery located in the lowlands of the central part of the Bohemian distribution of the CWC. The left-sided burial no. 10 contained an extraordinarily large number of drilled teeth and shell beads (probably the largest within the whole archaeological culture) arranged in strips (probably fixed to clothing). The finds represent at least 412 whole teeth from at least 78 dogs, and 4000-5000 shell beads, in addition two large solar discs of *Margaritifera auricularia* [1,2]. Two dog teeth from this collection, specifically 2x *incisivus 3 superior dexter*, were analysed here. The same anatomical position guarantees that the teeth originate from two different dog individuals.

#### Germany

##### *Bürgermeister-Ulrich-Straße 100, Augsburg, Germany*

The site is located in the south of today’s city of Augsburg, Bavaria (48.32351°N 10.89306°E) just a few hundred meters south of the Bell Beaker cemetery of “Augsburg – Universitäts- Sportgelände”. Large scale archaeological rescue excavations were carried out in 1986 until 2002 within the industrial park leading to the discovery of Early Bronze Age longhouses and several probably contemporaneous pits containing ceramic materials as well as animal bones, which are part of this study. Alongside these settlement evidences, undated ring ditches and Late Neolithic burials were discovered. The oldest burials are two graves situated next to each other attributed to the Corded Ware Complex. In total twelve Bell Beaker burials were found distributed in three separate groups. Graves 1-8 were found in the northwest of the excavation site, grave 9-11 further south next to the Corded Ware burials, but separated by approximately 10-15 meters and grave 12 was found around 120 meters northeast of them. The latter was constructed with a ring ditch. From the small faunal assemblage, two specimens have been selected for ancient DNA sampling: This includes a thoracic vertebra of a small domestic ruminant showing exostoses at the caudal surface of the Processus thoracalis and a cattle metacarpal with a possible exostosis on the dorsal side of the proximal diaphysis, even if this feature is not clearly visible due to strong bone surface erosion.

##### *Gewerbegebiet Nord, Kleinaitingen, Germany*

The site of Kleinaitingen – Gewerbegebiet Nord, Bavaria (48.22281°N 10.84499°E) is located ca. 16 km south of today's city of Augsburg on the upper gravel terrace of the Lech River. Apart from some prehistoric (mostly Late Bronze Age settlement) features, a large Early Bronze Age cemetery was documented between 2012 and 2014 by members of the local history and archaeology society (Arbeitskreis Augsburg Süd). It was almost entirely excavated; only the southern limit is still covered by a modern pathway. In total 63 graves with 72 individuals could be documented, most of them inhumations, making it one of the largest Early Bronze Age cemeteries in southern Germany [13]. All individuals/inhumations were buried in a crouched position with typical gender-specific orientation and body placement of the Bell Beaker and Early Bronze Age type. The radiocarbon dates span from 2116 until 1514 cal BCE, with the majority of the dates falling between 1900 and 1700 cal BCE. Animal bones were a part of the grave-good assemblages. For this study animal bones from four graves (nr. 30, 45, 52 and 62) as well as two pieces from a contemporary ring ditch (feat. 13) were sampled. Since due to the burial rites characteristic for the Early Bronze Age in the northern Alpine foreland, the faunal record in grave contexts is largely limited to teeth and mandibles. Thus, sampling focused on these skeleton elements. Five cattle teeth and one cattle mandible representing individuals of different slaughter age stages have been selected for this study.

##### *Obere Kreuzstraße, Königsbrunn, Germany*

The site of Königsbrunn – Obere Kreuzstraße is located ca. 10 km south of today's city of Augsburg, Bavaria (48.26656°N 10.87857°E) on the upper gravel terrace of the Lech River at the western edge of the town of Königsbrunn. It was excavated during preconstruction works in 2007 in its full extent. All 48 grave pits of the Early Bronze Age cemetery were oriented NNE-SSW or shifted slightly clockwise and comprised five distinct groups of graves [13]. In total 50 inhumations could be documented with individuals buried in a crouched position in typical Bell Beaker and Early Bronze Age gender-specific body positioning. The radiocarbon dates range between 2136 and 1780 cal BCE. Most of the dates fall between 2040 and 1880 cal BCE. Animal bones, mostly cattle teeth, were a regular part of the grave good assemblages. For this study six upper or lower cattle molars from different graves as well as a mandible and teeth from a contemporary deposition of a young cattle within the northern grave group (feat. 215) were sampled.

##### *Postillionstraße, Haunstetten, Germany*

The site of Haunstetten - Postillionstraße is located ca. 7,5 km south of today's city centre of Augsburg, Bavaria (48.29573°N 10.89103°E) on the upper gravel terrace of the Lech River in the southern part of the district Haunstetten. The Early Bronze Age cemetery was excavated in 1992 by the city archaeology of Augsburg before housing estates were built. Parts of the site could not be documented, but large areas of the Early Bronze Age cemetery were excavated under regular circumstances. Apart from the cemetery a small ditch and a horse burial from the Roman period were found. The Early Bronze Age cemetery contained at least 40 graves with 41 individuals [13]. All individuals were inhumations and buried in accordance with the gender-specific orientation and body positioning of the Bell Beaker and Early Bronze Age type. Five graves (feat. 44, 50, 65, 137 and 111) were accompanied by long post alignments, making them outstanding burial monuments/individuals. Radiocarbon dates gathered from skeletal remains range between 2197 and 1772 cal BCE. Most of the dates fall between 2140 and 1880 cal BCE. Most of the animal remains sampled in this study stem from postholes of these alignments. Additionally, one sampled tooth comes from a ring ditch (feat. 49) surrounding an Early Bronze Age burials (feat. 50), another tooth was part of the grave good assemblage of one of the burials (feat. 99). The faunal assemblage exclusively consists of cattle remains, of which five molars and two mandibles have been included in this study.

##### *Universitätsstraße, Augsburg, Germany*

The site of Augsburg - Universitätsstraße is located in the middle of the university campus in the south of today's city of Augsburg, Bavaria (48.33262°N 10.89402°E). Besides various prehistoric archaeological features a large pit with a substantial amount of Early Bronze Age pottery accompanied by animal bones was excavated (feat. 43). The pottery can be dated roughly around 1800-1600 BC. Five animal bones studied here stem from this feature, another from a most likely Early Bronze Age slit-pit. The faunal assemblage consists of only seven identified specimens, representing the livestock species characteristic for the Early Bronze Age economy of the Northern Alpine foreland: cattle (n=1), sheep (n=1), sheep/goat (n=2), and pig (n=3). From this assemblage, we selected a Phalanx proximalis (cattle), a Phalanx medialis (sheep) and a scapula (pig) for sampling.

##### *Unterer Talweg 85, Haunstetten, Germany*

The site of Haunstetten – Unterer Talweg 85 (today street nr. 49) is located ca. 4,5 km south of today's city centre of Augsburg, Bavaria (48.31966°N 10.89153°E). This large-scale excavation took place in 1999 and uncovered remains of Early and Late Bronze Age settlement remains as well as burials with inhumations from the Bell Beaker Period and Early Bronze Age [13]. The Bell Beaker graves have been found in two concentrations, ca. 170 metres apart from each other. The northern Group contained five single burials oriented N-S. The radiocarbon dates range from 2465–2143 cal BCE. The two single graves of the southern group could be contemporaneous to the burials of the northern group (2456–2141 cal BCE). Almost in the middle of both Bell Beaker burial groups, but ca. 70 metres to the east, a small part of a presumably larger Early Bronze Age cemetery was unearthed with radiocarbon dates spanning between 2025-1885 cal BC. West of these Late Neolithic and Early Bronze Age burials remains of a large Early Bronze Age settlement were uncovered with at least seven long houses accompanied by smaller storage buildings. On the western side of the excavated area at least two Late Bronze Age timber buildings could be identified as well as several La Tène Period (Late Iron Age) pits with human remains. Faunal remains include a total of 24 identified specimens, distributed among cattle (n=12), sheep (n=1), sheep or goat (n=2), pig (n=7), red deer (n=1) and beaver (n=1). Of these, 18 have been selected for sampling (several with multiple subsamples).

##### *Unterer Talweg 111, Haunstetten, Germany*

The site of Haunstetten – Unterer Talweg 111 is located ca. 5 km south of today's city centre of Augsburg, Bavaria (48.31421°N 10.89007°E). Four neighbouring plots were archaeologically investigated due to the construction of three supermarkets, among other things (Unterer Talweg 109-115). In the northern area, a circular ditch and several Late Bronze Age cremation burials came to light. The cremation graves extended as far as the plot at Unterer Talweg 113. At least seven Early Bronze Age buildings were uncovered in the area of Unterer Talweg 111, 113 and 115. Ten Early Bronze Age burials were found to the north-east of the settlement (Unterer Talweg 111) which can be dated to the middle phases of the Early Bronze Age (ca. 2050-1800 BC) [13]. The sampled animal bone material comprises a sheep's mandibular and a tooth row without alveolar bone of the same species.

#### Moldova

##### *Petreni, Moldova*

The settlement of Petreni is located on a hill plateau within the hilly landscape of the Bălţi Steppe. Belonging to the Cucuteni-Tripolye-Complex, where settlements, especially in the first half of the 4th Millennium BC, show the tendency to form huge agglomerations of houses, which occupy very large spaces of up to 320 ha. The houses were built and appear to have been deliberately burnt within a rather short period of less than 200-300 years. Well-preserved zooarchaeological remains were recovered during the excavations led by Dr R. Uhl. A total of 3146 bones were identified and taxonomically assigned by N. Benecke and M. Hochmuth (unpublished). Cattle were the predominant species (Number of identified specimens (NISP)=1515) while sheep/goat and pigs represented 34% and 10.6% respectively of the identified material. For study, 30 samples were selected by R. Uhl, R.E. Gillis and A.K.W. Runge, from secure contexts.

#### Poland

##### *Giecz 10, Poland*

This site is located in central Poland in the Wielkopolska region (52°19'9.00" N 17°22'14.99" E) and was discovered by Józef Kostrzewski in 1928 [14,15]. The first excavation was carried out in 1948 by Bogdan Kostrzewski [16]. Giecz was one of the most important centers in the formation of the Polish state during the early Middle Ages. Historical records and particularly archaeological research indicate that there was a stronghold of great political, economic, and military importance located at Giecz from the second half of the ninth century to the thirteenth or fourteenth century. Numerous settlements and cemeteries were located around this stronghold. In recent years, archaeological research at Site 10 has revealed one of these settlements. Materials collected from numerous contexts, including pottery, clay, metal, and stone artefacts, as well as jewelry and bones of both humans and animals, are sources of data on the chronology, demography, and everyday life of the site, and can be employed as indicators of formational and post-depositional processes. In this way we know, mainly from the study of pottery, that the settlement at Site 10 existed between the ninth and tenth centuries, and was therefore contemporaneous with the oldest pre-state development phase of the Giecz stronghold—which highlights its importance in research. A total of 62,466 animal bones were obtained from the archaeological contexts of Site 10 during archaeological investigations in 2014–2019 and 2021–2023, some of which have undergone zooarchaeological and taphonomic studies by Pawłowska [17–20]. Overall, the proportion of specimens with pathological changes did not exceed 4% [19–22]. Changes such as arthropathies, inflammatory diseases, diseases connected to the environment, dental anomalies, oral pathology, and traumatic lesions were found. In this study, fifteen subsamples representing different skeletal elements were selected for DNA analysis (**Table S1**); these come from cattle, pigs, sheep or goats, and red deer.

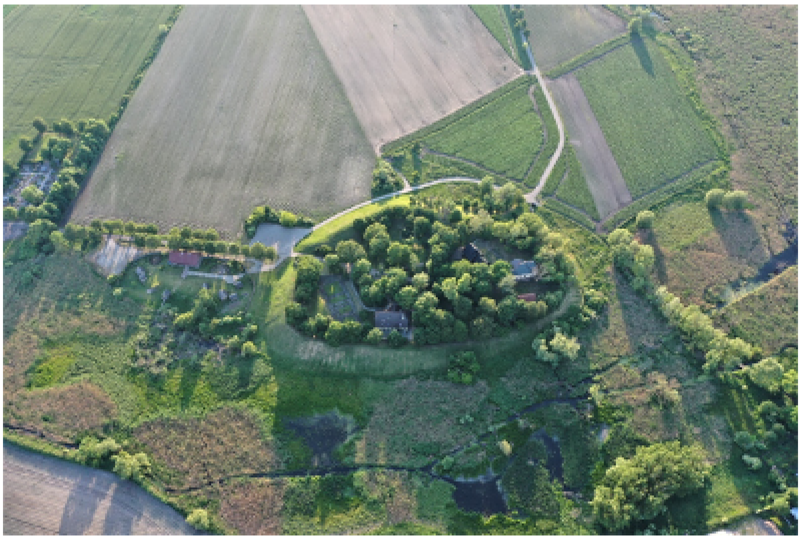

Photo of the Giecz site, view of the stronghold. Photo Marcin Krzeptowski.

##### *Zdrojówka, Poland*

This archeological site is located in central Poland in the Kujawy region (52º26'16" N 18º44'50" E). It was discovered in 1957, when a human grave and a neighboring double burial of cattle, belonging to the Globular Amphora Culture, was uncovered [23,24]. The west-to-east oriented grave was surrounded by stones and clay. Blades were reported from this context, as well as other artifacts such as amphorae [23,25]. The burial includes examples of ritual patterns of animal deposits equipped with pots [26]. Among the specimens, a tooth was selected for study: this specimen does not show evidence of pathology.

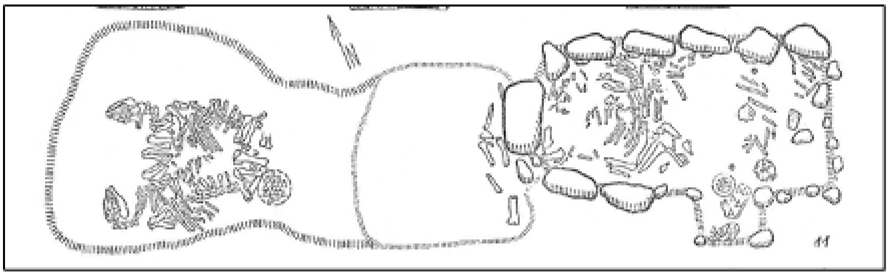

The Zdrojówka site: grave with human remains (on the right) and double cattle burial (on the left). (Wiślański, 1963)

##### *Gdańsk, Św. Wojciecha site, Poland*

This site is located in the city of Gdańsk in Pomerania in the northern part of Poland (54º16'3.63" N 18º39'5.61" E). Archaeological research was conducted here in the 1950s. The archaeological contexts of the site have been dated to the ninth to thirteenth centuries [27]. The animal remains were studied by Kubasiewicz [28], though specimen pathology was not then reported. However, a pathological study was carried out by Pawłowska in 2021. Among the specimens with pathological changes, a second phalanx of cattle was selected for study: this specimen shows a lytic lesion within the shaft.

##### *Site 63, Krzczonowice, Poland*

Archaeological site no. 63 at Krzczonowice (50°51'56"N 21°29'43"E), in Świętokrzyskie province, Poland, is a part of the larger excavation area within the Archaeological Museum and Krzemionki Reserve (UNESCO), which was initiated in the 1930s [29–31]. The explorations at Krzczonowice, which began in 2004, are located in the middle and lower basin of the Kamienna river in the northern part of the Sandomierz Upland. Indirect chronology has dated this settlement to the Globular Amphora Culture. Direct radiocarbon dating has shown that a bone fragment from the site comes from 2476–2293 cal BC (4425–4242 cal BP). Special attention was paid to object no. 33, which was identified as a sacrificial pit with an almost complete bovine skeleton deposit. The depth of the oval pit is ca. 180 cm and it contains the skeletal remains in partly anatomical order [31]. The total number of bone fragments (TNF) was 302 and species identification was possible in the case of 78.15% items (NISP = 236). Over 85% of the NISP was skeletal material coming from a taurine cattle individual. Other species include pig, sheep/goat, horse, lynx, and humans. Before deposition, the cattle cadaver was probably partitioned. The thoracic, lumbar, sacral, and caudal part of the vertebral column, together with the right pelvic limb, was unearthed as articulated laying on the left flank. The position of the cervical vertebra and the mandible suggest that the head and neck were bent caudally and positioned on the right side of the animal trunk. The skull was unhorned. The bones of the left pelvis and of both thoracic limbs were not in anatomical position. The only visible pathological feature was noted within the greater trochanter of the right femur, as a massive bone tissue loss process that led to the formation of a concave area (cortical and spongy bone destructive and lytic process). Further paleopathological analysis may be able to identify the potential cause of the disease. Genetic analysis was carried out using bone material from the lesion location.

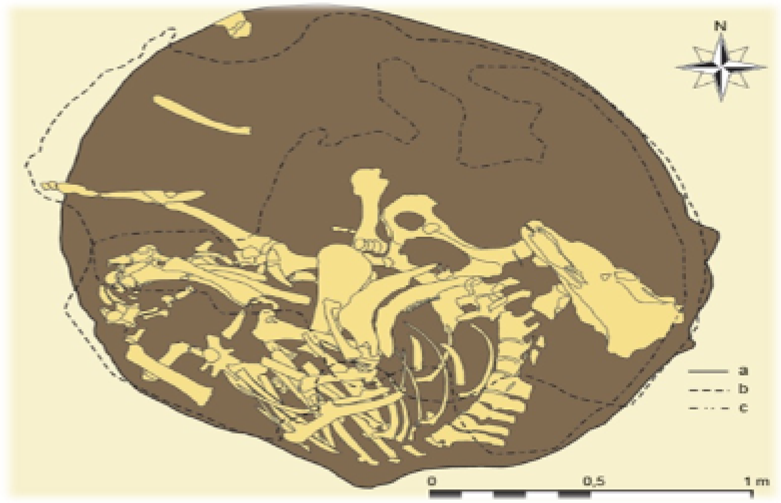


The Krzczonowice 63 site, burial of cattle (pit no. 33). Photo: Artur Jedynak

##### *Site 154, Inowrocław, Poland*

This site is located in the Kujawy region in Poland (52º47'42.87" N 18º15'34.42" E). The archaeological rescue excavation at Site 154, Inowrocław, was conducted in 2009 by Marcin Woźniak. The stratigraphy of the site, along with the archeological context, was established using ceramics found to be from the Bronze Age (VEB–HaC: 8th-7th century BC). During the excavation, animal remains were recovered from the settlement complex, of which some assemblages were studied [32]. Pathological lesions were found on twenty specimens. In the present study, seventeen subsamples representing a range of skeletal elements were selected for DNA analysis, as they showed inflammatory diseases (**Table S2**). These specimens represent horses, aurochs, cattle, pigs, sheep or goats, and dogs.

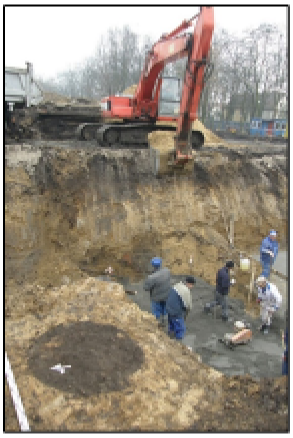

The Inowrocław 154 site, excavation area. Photo: Marcin Woźniak

##### *Moszna Wieś, Poland*

The site is located in the Mazowsze region in northeast Poland (52º10'51.28" N 20º44'50.18" E). Archaeological research was conducted at the site, leading to the discovery of a settlement with features of the Jastorf culture; dating to the younger pre-Roman period in Moszna Wieś was based solely on the analysis of ceramic material. Animal bones were also recovered during this work (NISP = 255). The assemblage was studied taxonomically, demographically, and taphonomically [33] and three specimens with pathological changes were found. Two subsamples, both mammalian long bones, were selected for the DNA analysis, as they showed inflammatory diseases (**Table S2**). Their state of preservation did not allow for species determination, but it was possible to assign them to the cattle size class.

##### *Izdebno Kościelne, Poland*

This site is located in the Mazowsze region in northeast Poland (52º08'15.52" N 20º31'18.67" E). The site was discovered in 1971 by Woyda, and archaeological research was conducted in 1976–1978 by Górna and later Nowakowski, and then again in 2008-2009 by Rozen and later Piłatowski. The archaeological contexts and recovered materials were dated to the Paleolithic, Neolithic, early Bronze Age, early Iron Age, pre-Roman period, Roman period, and early phase of the migration period, as well as late medieval and modern times [34]. Archaeological structures were uncovered and bulk materials were obtained, including more than 6000 animal bones. These collections have been investigated in terms of taxonomy, demographics, and taphonomy [35]. 36 specimens with pathological changes were found. Two subsamples were selected for the present DNA analysis (**Table S2**); one each from the contexts dated to older and younger pre-Roman period, as well as the late Roman period. These elements both originated from cattle.

#### Romania

##### *Pietrele-Măgura Gorgana (Pietrele), Romania*

The site is situated close to the Danube river in Romania (44.068869°N, 26.159005°E) and is recognized as a nine-meter-high tell settlement in the archeological record. Excavations have led to the discovery of cultural layers that are dated to 4550 – 4250 BCE and yielded comprehensive assemblages of animal remains, which were used to infer the economic basis and local environmental conditions [36,37]. From a chronological point of view, the finds can be assigned roughly to two phases in time, namely the Late Neolithic Boian culture (the outer settlement) and the Copper Age Gumelnita culture (the tell) [38]. Animal remains obtained from these contexts and studied by Benecke [36–38]. The Boian settlement in Pietrele is mainly typical of an agrarian-oriented food economy focused primarily on cattle (70%), followed by pigs (15%) and small ruminants (sheep and goats; 15%). The proportion of wild boar and red deer did not exceed 8%. In contrast, husbandry from the subsequent Gumelnita settlement attests to husbandry mainly of pigs (56%), followed by cattle (33%) and sheep and goats (11%) [38]. K. Pawłowska conducted a study in 2022–2023 to select specimens with pathological changes for this study. 83 skeletal elements (AZP-102–AZP-184) were selected for DNA analysis; these showed inflammatory diseases, dental anomalies, oral pathology, and inflammation associated with arthropathy. These specimens represent cattle, pigs, sheep, goat, sheep or goats, dogs, aurochs, wild pig, red deer and beaver (**Table S2**).

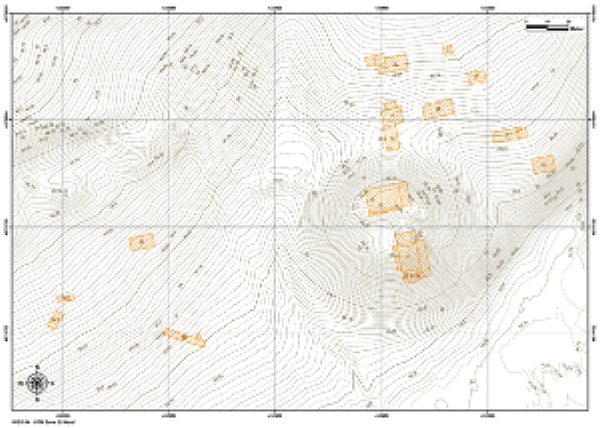

Image of the Pietrele site, with the tell and its immediate surroundings and excavation areas marked (D. Nowacki from Benecke et al., [38]).

#### Russia

##### *Ipatovo 3, North Caucasus, Russia*

The so-called 'Great Barrow' of Ipatovo, kurgan 2, is one of the most important Bronze Age sites in the steppes north of the Caucasus. It is located on the outskirts of the city of Ipatovo, about 120 km northwest of Stavropol, Russia. The mound is one of three larger mounds and part of a long line of smaller ones. It is situated in the valley of the Kalaus River, which rises on the northern slopes of the Caucasus Mountains and flows into the Manych River to the north that is an important north-south connection on the eastern edge of the Stavropol Upland between the Caucasus mountains and the steppes.

Kurgan 2 was one of the largest burial mounds in the area, measuring 7 m in height and over 100 m in diameter, and contained three different phases of Bronze Age occupation and a later reuse during the Sarmatian epoch. The mound was excavated in 1998/1999 over a period of one and a half years as part of a rescue excavation by the heritage organization ‘Nasledie’ in Stavropol, directed by A. B. Belinskiy [39]. The Bronze Age graves are part of bioarchaeological studies including isotopic analysis, paleogenetics and palaeoproteomics [40–44].

##### *Komsomolec 1-Marfa, North Caucasus, Russia*

The huge burial mound Komsomolec 1-Marfa was a single standing mound on the right river terrace of the Zolka creek, located 7,2 km west of the north-western outskirts of the city of Novopalovsk, Stavropol region, Russia. The area is today a piedmont steppe environment but was probably more densely forested in the past. The watershed along the river is covered with former barrows, now chiefly ploughed out. The site was excavated in a joint expedition of the local heritage organization ‘Nasledie’ and the Eurasia department of the German Archaeological institute, directed by A.B. Belinskiy and A. Nagler. Beside the mound a circular ditch enclosure is visible on aerial and satellite images, as well as in a magnetometry survey conducted in 2009 and 2011. The mounds height was 8,5 m, it had an oval shape, diameter along west-east axis was 62 m, along north-south axis 52 m. The mound was built in several consecutive years and altogether 60 graves have been excavated in the mound. The initial mound dates to the Early Bronze Age, the local late Maykop epoch, yet the central grave was destroyed. Nine more graves have been dated Maykop subsequent graves have been excavated as well, 48 date to different cultures of the Middle Bronze Age, 7 are of Late Bronze/Early Iron Age date, one Sarmatian and three modern individuals were found. Beside the burials a ditch with horse skeletons was opened, probably dating to Sarmatian times. Additionally, ritual areas with human skeletons were discovered beside the mound, dating to the Early Iron Age and the medieval period.

The site is prepared for a full publication and is central in bioarchaeological studies including isotopic analysis, paleogenetics and palaeoproteomics [40,43,44].

Grave 53, from where the studies sample originate, is a single inhumation of the Catacomb culture (2800-2600 calBC). The skeleton was found in a catacomb grave in a crouched position on the left side. The floor sprinkled with chalk and below the individual organic brown matter was found. Grave gifts are a bronze knife and two vessels and animal bones. The incisor of a caprid was studied.

##### *Marinskaya 5, North Caucasus, Russia*

Marinskaya 5 was another huge burial mound, located near the village Marinskaya, Stavropol region, Russia. The single-standing burial mound was excavated in 2009 by a team of the Lomonosov Moscow State University, the Institute of Archaeology RAS and the heritage organization 'Nasledie', directed by A.R. Kantorovich and V.E. Maslov. The kurgan is situated on the high terrace of the river Kura. The mound was slightly oval with a diameter of 34 to 40 m and 4,3 m high. At a distance of 10-16 m a ditch of 1,5 m depth surrounded the mound, which was detected on aerial images and excavated later. In the central part of the mound, the excavations uncovered three mound-shells phases dating to the Early Bronze Age Maykop epoch and a fourth one that was constructed during the Middle Bronze Age. A sequence of six burials date to the Maykop period, 17 Middle Bronze Age graves are associated with the local North Caucasian culture and one is a Catacomb grave culture grave and 3 date to the final Middle Bronze Age. In the Sarmatian period 6 graves and several ritual complexes were added to the older mound.

In several graves, among them late Maykop grave 25 and the North Caucasus graves 19, 23, 30/30a, paired cattle skulls were found. They place this site among one of the earliest where the use of cattle as draught animals is documented in a chronological sequence.

The site is partly published and central to a multidisciplinary study including stable isotope analysis, physical anthropology and paleogenetic investigations [40,43–46]

*Ransyrt 1, North Caucasus, Russia*

Ransyrt-1 is a wall system consisting of four semicircular ramparts with a diameter of about 300 m near the modern city of Kislovodsk in the region of the Caucasian Mineral Waters, Russia. The site is located at an altitude of about 1,800 m above sea level on the edge of a high plateau above the Podkumok River, at one of the most important pass roads in the central part of the North Caucasus. Three semicircular ramparts enclose a central area, which is itself surrounded by a wall. A magnetometer survey indicated intensive use of the site. Joint excavations in 2015 by heritage organization 'Nasledie' and the Eurasia department of the German Archaeological institute, directed by A.B. Belinskiy and S. Reinhold revealed stone structures and sunken rooms, for which the finds and features suggest ritual use [47]. All features date to the start of the Late Bronze Age (1850-1600/1500 cal BCE).

Four main phases of use were identified and dated, the first of which is the active use of the structures and during which an exceptionally large amount of animal bones and ceramics were left at the site. Presumably, they are the remains of ritual feasts or sacrifices. An analysis of the animal bones supports the thesis of selective, ritual use and suggests that those consumed in Ransyrt 1 were not kept in the surrounding area but came to the settlement from outside [48].

The cow tooth AZP-013 examined here comes from the Late Bronze Age stock of animal bones, while the souslik mandibula AZP-014 was excavated from the Late Bronze Age layers but might equally be a later intrusion.

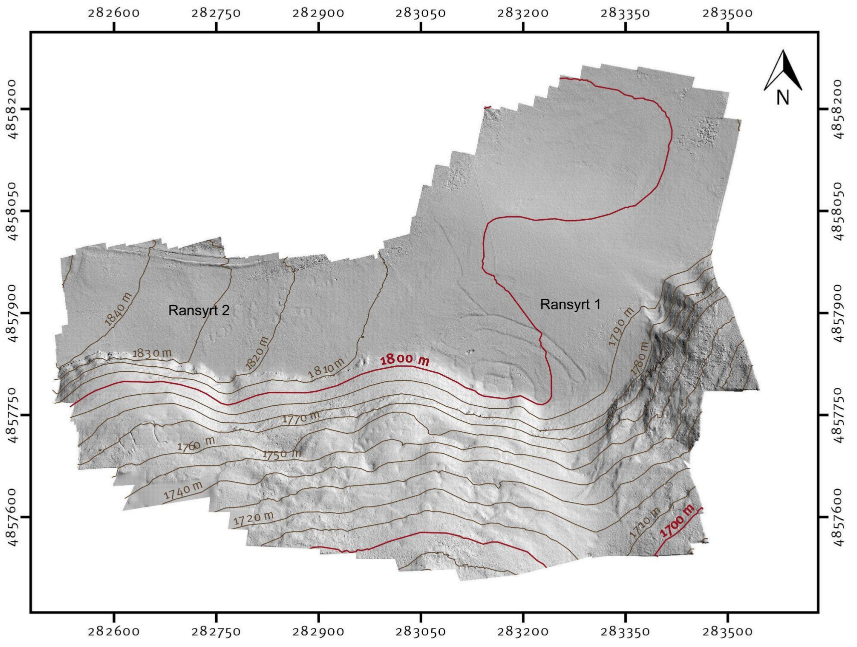


The sites of Ransyrt 1 and 2 situated at the edge of Podkumok river gorge © S. Rienhold, D. Mariaschk, DAI Eurasia department.

*Ransyrt 2, North Caucasus, Russia*

The second site, Ransyrt 2, is about 200-300 m west of the outermost rampart of Ransyrt 1. The site consists of a linear cluster of small stone socketed. Surface collections revealed Late Bronze Age material, but no excavations have been carried out so far.

The rodent bones examined come from surface collections and may stem from the Bronze Age cultural layers or are modern intrusions.

*Sharakhalsun 3, North Caucasus, Russia*

Sharakhalsun 3 is an archaeological site in the Manych Steppe, about 50 km southeast of the town of Divnoe, Stavropol region, Russia. It is one of several long rows of burial mounds running parallel to the Kalaus River. Many similar rows of burial mounds are located nearby, forming a unique burial landscape.

In 2001, 12 barrows were documented and five of them excavated in the course of rescue excavations by the heritage organization 'Nasledie', directed by A. V. Yakovloev.

Mound 1, from which the studied dog remains originate, dates to the Middle Bronze Ages' late Catacomb Culture (2400-2200 cal. BCE). The tooth itself was discovered in the mound fill and can either date to the Bronze Age or is a later intrusion.

*Sharakhalsun 6, North Caucasus, Russia*

Sharakhalsun 6 is another line of burial mounds in the Manych Steppe, about 50 km southeast of the town of Divnoe, Stavropol region, Russia. It runs parallel to the former site and equally to the Kalaus River. In 2001 during rescue excavations, 16 mounds were excavated by the heritage organization 'Nasledie', directed by A. V. Yakovloev. It is part of a bioarchaeological study including stable isotope analysis and paleogenetics [40,44].

The studied cattle tooth originate from mound 3 which is dated to the late Catacomb Culture (2400-2200 cal. BCE). The animal bones were found in a ritual context in the mound shell. It is 'animal 1' in the archaeological documentation.

*Stoderovskoe 1, North Caucasus, Russia*

The site of Stoderovskoe is a settlement dating to the Early Bronze Age Maykop epoch. It is located near the village Stoderovskaya in Stavropol region, Russia, about 13 km east of the city of Mozdok. The site is located in the valley of the Terek river, on its left bank.

During rescue excavations in neighbouring burial mounds in 2008 several pits with large amounts of pottery and animal bones and animal bone assemblages were excavated by an expedition of the heritage organization 'Nasledie' under the license of A. V. Lychagin. The studied cattle and pig bones stem from one of the pits and have also been investigated for stable isotope analysis and are in preparation for publication.

##### *Tomatniy 1, North Caucasus, Russia*

The burial mounds of Tomatniy 1 are located in the vicinity of the city of Pyatigorsk, Stavropol region, Russia. For the baselines of a stable isotope study, animal bones were collected in the storage of the heritage organization 'Nasledie' from mound 1, grave 8 that dates to the Bronze Age. No further information is available.

#### Turkmenistan

##### *Monjukli Depe, Turkmenistan*

The small mound of Monjukli Depe in the foothills of the Kopet Dag in south Turkmenistan (36.8484° N, 60.4180° E) was occupied in the sixth millennium BCE during the Neolithic Jeitun period and, after a long hiatus, reoccupied in the Early Aeneolithic (Chalcolithic) in the fifth millennium BCE. Initially discovered and excavated in Soviet times [49], fieldwork at the site was resumed from 2010 to 2014 by a team from the Freie Universität Berlin. The Neolithic levels excavated in only small proportions were characterized by diffuse layers with only little architecture, containing variable densities of artifacts. The extensively investigated Aeneolithic levels exposed substantial portions of superimposed and well-preserved mudbrick architecture [50,51]. All samples are from features of the Aeneolithic occupation. The excavations at Monjukli Depe have uncovered an extensive collection of approximately 53,000 animal bones. The taxonomic identification of the material was conducted by M. Hochmuth and N. Benecke. They consist predominantly of remains from domesticated caprines (>90%) both in the Neolithic and Aeneolithic assemblages, with sheep outnumbering goats [52–55]. A markedly smaller number of domesticated cattle were recovered. Dog bones complete the spectrum of domesticated animal taxa. The inhabitants of Monjukli Depe focused almost exclusively on the rearing of small ruminants. Wild fauna accounts for less than 5% of the identified remains, with species including onager, gazelle, wild sheep, and foxes, indicating that hunting played a very minor economic role for the inhabitants. Multi-isotope analysis of tooth enamel (carbon, oxygen, strontium) and bone collagen (carbon, nitrogen) suggests substantial dietary variability among caprines, with evidence of $C_4$ plants contributing to the diet of part of the herd on both seasonal and long-term scales [55,56]. This indicates herding practices that utilized the micro-variability of the local landscape, with sheep and goats kept in small flocks across ecologically diverse areas close to the settlement.

#### Uzbekistan

##### *Tilla Bulak, Uzbekistan*

This site is situated near the border tripoint of Uzbekistan, Turkmenistan, and Afghanistan, in ancient Bactria (UTM 42S 306120 m E / 4176000 m N ), about 850m above sea level. The settlement here developed in the early phase of the Late Bronze Age, which can be correlated with the Sapalli Culture—that is, a local variant of the Southern–Central Asian Namazga Cultures. The site extends over an area of about 0.4 ha and is dated to the twentieth to nineteenth centuries BC. Exploration in this area has been carried out by the Institute of Near Eastern Archaeology, Ludwig Maximilian University of Munich, by the Tocharistan expedition of the National Institute of Fine Arts in Tashkent, and by the University of Termez. The excavations on site were conducted from 2007 to 2010, uncovering almost 40% of the examined settlement area [57,58]. Animal bones (NISP = 10,869) were recovered from the archaeological contexts of the two main phases of the site. Sheep and goats dominate the assemblage (92%), with cattle representing 5% of the assemblage, while the proportion of pigs was negligible [59]. The original analysis recorded no pathologies. More recently, K. Pawlowska and R. E. Gillis have recorded a high frequency of pathologies, particularly within sheep/goat remains. Of these, 105 skeletal elements were selected for study, as they showed inflammatory diseases, dental anomalies, oral pathology, inflammation associated with arthropathy, and traumatic lesions. These specimens represent horses, cattle, sheep, goat, sheep/goats, dogs, fox and gazelle (**Table S2**). A fragment of a cattle rib (AZP-285) displays a lesion that could be evidence of a neoplastic, tumorous, or inflammatory disease such as, osteolytic cancer or tuberculosis, respectively.

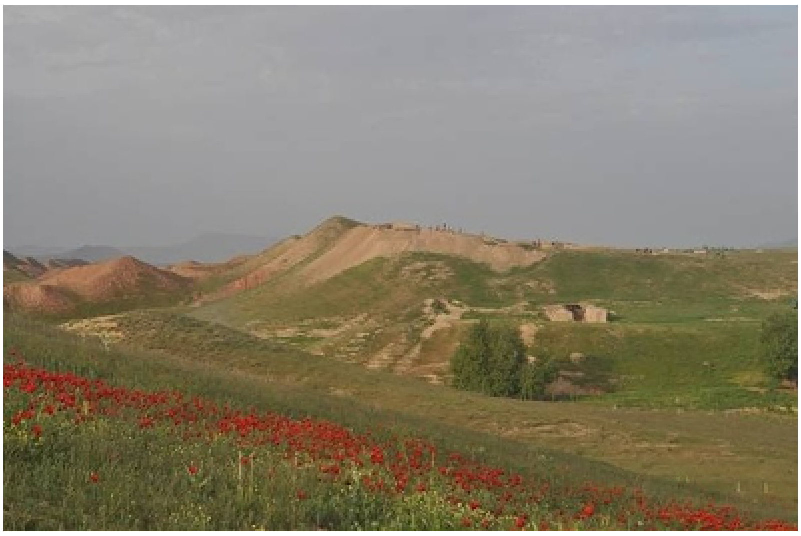

Photo of the Tilla Bulak site (by K. Kaniuth).
